# Pleiotropic Regulatory Locus1 maintains actin microfilament integrity and concomitant cellular homeostasis facilitating root development in Arabidopsis

**DOI:** 10.1101/2023.02.28.530538

**Authors:** Chi Wang, Xing Wang, Zhenbiao Yang, Xiaowei Gao

## Abstract

Cell functions are based on integrity of actin filaments. The Actin cytoskeleton is typically the target but also the source of signals. An evolutionarily conserved WD-40 protein PRL1 (Pleiotropic Regulatory Locus1) in Arabidopsis was investigated with multilayer functions in development, innate immunity, alternative splicing activation, transcription regulation, genome maintenance, ubiquitination-based protein turnover et al., but the underlying mechanisms are undefined. Here, we show PRL1 maintains actin integrity and concomitant cellular homeostasis. To explore causes for developmental root defect, we found depolymerization of cortical actin cytoskeleton and ROS imbalance in *prl1* mutant. Further, we revealed that actin de-polymerization was the fundamental cause and dominant to ROS imbalance (H_2_O_2_ and O_2_^·–^) for retarded root of *prl1*; *NAC085* was up-regulated by and cooperated with actin depolymerization to mediate to stele cell death. Moreover, we revealed stress-related differentially expressed genes and alternative splicing defects were mutually independent and were responses to actin depolymerization in *prl1*. Our work ravels out cause-effect relationships between actin configuration and downstream hierarchical signals and explores underlying mechanism for functions of *PRL1*.

## Introduction

PRL1 (Pleiotropic Regulatory Locus1) was explored with multiple biological and cellular functions by analyzing various defects in its null mutants (Baruah et al., 2009; Nemeth et al., 1998; Palma et al., 2007; Zhang et al., 2014). In Arabidopsis, an allelic *prl1* mutant was first isolated for finding sugar signaling component and found with multiple defects in root development, hypocotyl elongation, and enhanced hypersensitivities to various phytohormones marked by their developmental phenotypes (Nemeth et al., 1998). Later, PRL1 was also found to be a core component in MAC/NTC complex and responsible for plant innate immunity (Monaghan et al., 2009; Palma et al., 2007). Additional experiments support that PRL1 is involved in activation of spliceosome complex, microRNA processing, genome maintenance (Ajuh et al., 2001; Jia et al., 2017; Koncz et al., 2012; Li et al., 2018; Zhang et al., 2014), and protein turnover (Bhalerao et al., 1999; Farras et al., 2001; Lee et al., 2008). So far, the links between its biological functions and molecular, biochemical functions of *PRL1* were not established, that is to say the mechanisms underlying its multiple biological functions are undetermined.

A striking characteristic of *prl1* mutant is short primary root (Ji et al., 2015; Nemeth et al., 1998; Wang et al., 2021), but the mechanism underlying its developmental defect is uncovered. At cellular levels, root length is directly determined by combination of controlled cell division in meristem (MZ) zone and regulated anisotropic cell expansion in the elongation zone (EZ) (Beemster and Baskin, 1998). As we known, ROS (Reactive Oxygen Species) plays dual roles in regulation cell actions. In developmentally and environmentally favorable conditions, ROS keeps balance and takes the instructional role of facilitating root development by controlling spatial transition from cell proliferation to cell differentiation which was recently revealed to be mediated by UPBEAT1-controlled ROS distribution (Tsukagoshi et al., 2010); another, in response to encountered exogenous or endogenous stresses, ROS imbalance is generated, which takes series of completely detrimental roles in reducing MZ size by accelerating the proliferation-to-differentiation transition, preventing cell expansion by decreasing cellular membrane fluidity and increasing cell wall solidity and even damaging DNA and initiating resultant DDR (DNA Damage Response) including cell cycle arrest, cell death and growth inhibition (Baxter et al., 2014; Mittler et al., 2022b; Petrov et al., 2015). In *prl1* mutant, the previously published data showed the stress responsive genes were de-repressed, indicating intracellular stress signal were produced (Jia et al., 2017; Salchert et al., 1998). Did that mean endogenous stress was the cause of root developmental defect in *prl1*? Where was endogenous stress derived?

In both plants and animals, the cytoskeleton is essentially the basic and structural component for cell behaviors including cell divisions and cell expansion and is typically considered the target of signals (Blancaflor et al., 2006; Vaughn et al., 2011a). Actin Dynamic homeostasis is required for its normal functions. The dynamics of actin filament is controlled by two types of actin binding proteins (ABPs), one serving as actin filament polymerization factors and the other as actin filament de-polymerization factors, which are elaborately regulated by upstream signals to maintain dynamic actin homeostasis (Kadzik et al., 2020; Li et al., 2015). Given intracellular actin could form various configurations, it is important for cell to sense actin configuration and make feedback to its reorganization. But when dynamic actin cytoskeleton was out of homeostasis, what kind of signals would be recognized and transmitted by cells? Recent works presented that abnormal cortical actin structures would be recognized as intracellular dangerous signals (DAMPs) to stimulate ROS accumulation, DNA fragmentation and widespread transcriptional changes of defense/stress responsive genes (Brown, 2012; Desouza et al., 2012; Franklin-Tong and Gourlay, 2008; Leontovyčová et al., 2020; Smertenko and Franklin-Tong, 2011; Thomas and Franklin-Tong, 2004; Thomas et al., 2006). Together, actin cytoskeleton takes dual character as cellular scaffold and as signal.

In this study, we focus on exploring mechanisms for developmental defect in the primary root of *prl1* mutant and try to investigate general mechanisms for biological functions of *PRL1*. Through a series of cellular, pharmacological, genetic and molecular investigations, we found that cortical actin was depolymerized and revealed its de-polymerization was the fundamental cause for short root of the *prl1*; ROS imbalance was a type of stress response to actin de-polymerization; *ANAC085* was induced by and specifically cooperated with actin cytoskeleton de-polymerization to implement root stele cell death and the de-repression of stress/defense-related genes was also a secondary response to actin de-polymerization and independent of alternative splicing defects in *prl1*. Thus, it is revealed that the basic function of *PRL1* is to maintain actin integrity and concomitant cellular homeostasis. This result presents a completely new recognition of functions of *PRL1* and would inspire further investigations on molecular mechanisms for its functions.

## Result

### Mutation in *PRL1* leads to root growth retardation

Developmental root defect is a remarkable character of the *prl1* mutant, but the underlying cellular mechanism is still unknown. To reveal general mechanism underlying its multiple biological functions, we focused on revealing mechanism for root developmental defects in the *prl1*.

Firstly, we checked root length differences between WT and the *prl1*. We found the primary root of *prl1* was serious shorter in 6 DAG (Day After Germination) than WT and the primary root growth was quantitatively counted to be significantly inhibited in *prl1* over time (Figure 1, A and B). As root meristem was the source of cells supplying continuous root growth, we checked the root meristem size and corresponding meristem cell numbers of *prl1* to be smaller and less (Figure 1, C and D) indicating reduction of cell production or early differentiation or both. The length of the cells in counterpart root mature zones of *prl1* is shortened in contrast to WT (Figure S1A) indicating cell elongation is inhibited. Cell is the final functional target of signal and the basis of root development, we checked root cells statuses in WT and *prl1*. Using constructed *proCycB1;1-GUS prl1* double mutant, we observed GUS staining was almost vanished in *prl1* background (Figure 1F), indicating cell division activity was reduced in meristem zone compared to WT. Using local WOX5 expression to mark the location of QC, we found expanded distribution of pWOX5-droven GFP in *prl1*, indicating extension of QC identity to surrounding cells (Figure 1E) which also was strengthened by extended *QC25: GUS* distribution (Figure S1C). At normal conditions, QC keep constant lower division activity, those results implied QC was encountered endogenous stress (Heyman et al., 2014). Using PI staining, we found permeation of PI into stele cells above QC indicating existence of cell death (Fulcher and Sablowski, 2009) which also indicated existence of intracellular stress. By time-laps observation, we found stele cell death was not emerged until 3DAG and QC expansion was appeared after stele cell death (Figure S1B) suggesting cell death was genetically programmed. In addition to encountered intracellular stress, QC expansion might be the result of stele cell death which deprived the restricting signal for QC expansion.

**Figure 1.**
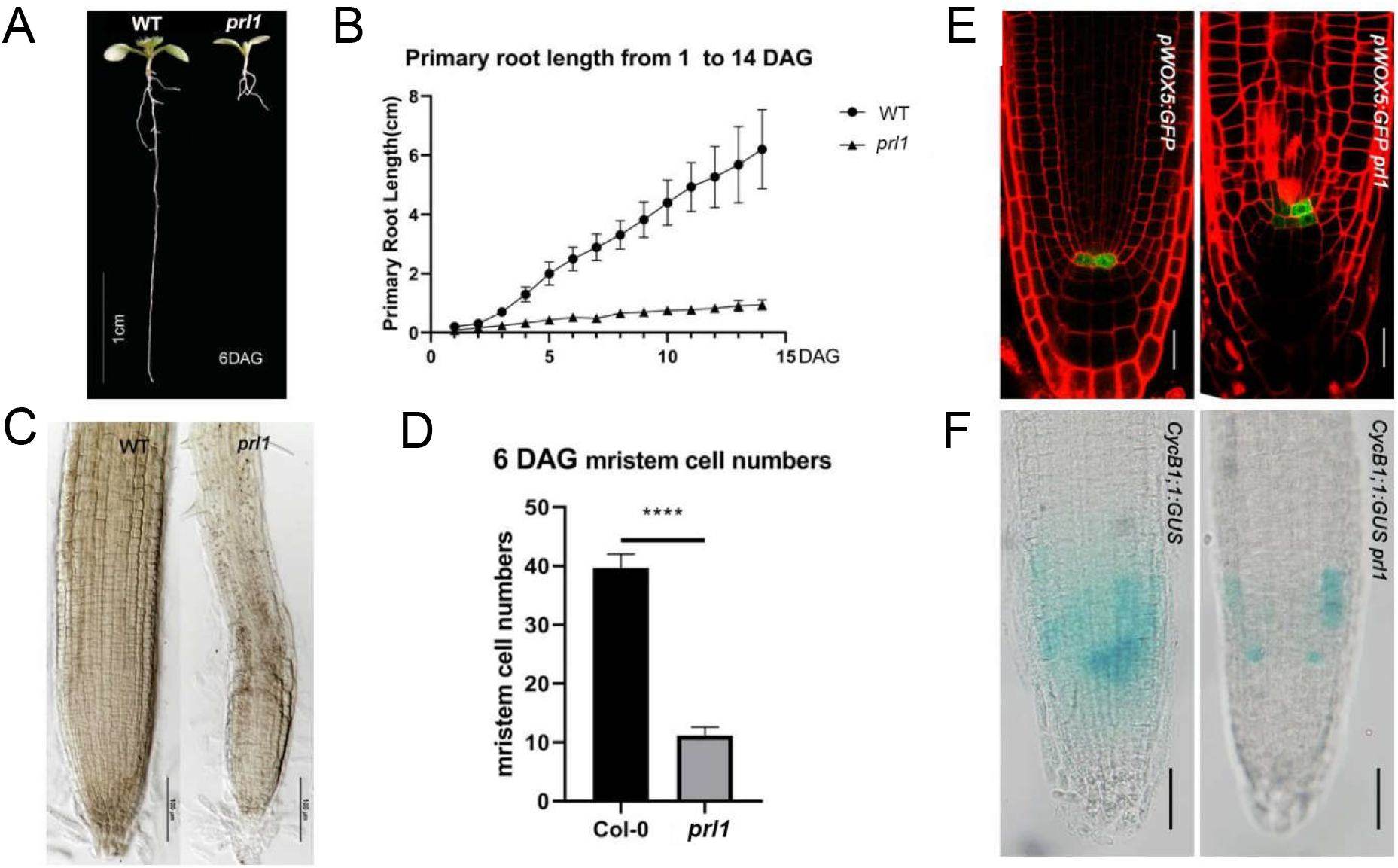
Null mutation of PRL1 retards primary root development. (A) Representative root images of WT and *prl1* in 6-DAG (day after germination) seedlings vertically cultured in ½ MS. Bar=1cm. (B) Quantification of primary root length of WT and *prl1* from 1 DAG to 15 DAG. Values are means ± SD of more than 50 primary root length. (C) Representative root meristem region of WT and *prl1* in 6-DAG seedlings. The size of root meristem region is typically defined from QC to distinctive cortical cell with double size to its neighboring QC-closer cell. Scale bar = 100 μm. (D) Quantification of cell numbers in meristem zones of WT and *prl1*. values are mean ± SD of more than 30 roots at 6 DAG. Asterisk indicates significant difference (****p<0.0001, t test). (E) *pWOX5:GFP* expression patterns and PI staining-displayed stele cell status in 6-DAG WT and *prl1*. Scale bar = 20 μm. (F) CyclinB1;1: GUS expression levels in 6-DAG WT and *prl1* roots. Scale Bar = 20 μm.

Taken together, cells statuses indicated *prl1* root cells were wholly disrupted, and according to general disruption of root cell behaviors such as cell cycle arrest, cell death and QC expansion, we predicted cells of *prl1* root were subjected to intracellular stresses.

### Mutation in *PRL1* leads to ROS imbalance which is enough to cause short root

ROS imbalance is always considered the intermediate product in response to encountered stresses (Jaspers and Kangasjärvi, 2010). To check changes of ROS in *prl1*, we used DAB and NBT solutions to stain roots of WT and the *prl1* at 6 DAG to detect H_2_O_2_ and O_2_^·–^ respectively (Zeng et al., 2017). We found in *prl1* the DAB staining intensity was stronger along almost the whole longitudinal primary root and invaded to site where meristem zone is normally localized in WT (Figure 2A), indicating increase of concentration and expansion of H_2_O_2_ and premature differentiation of root meristem cells. The observed increase of root hair density in *prl1* also indicated of H_2_O_2_ accumulation (Figure 2C). Along longitudinal axis of primary root, the NBT staining intensity was observed to be weaker and the NBT staining marked O_2_^·–^ distribution gradient in WT was disappeared in *prl1*, suggesting resettlement of O_2_^·–^in *prl1* root (Figure 2B). To test whether ROS imbalance was enough to hinder primary root length, we resettled ROS in WT utilizing chemicals to mimic those in *prl1*. As H_2_O_2_ and O_2_ ^· –^ levels and distribution were observed spatially antagonistic, we used two types of scavengers N-propyl gallate (PG) and Diphenyleneiodonium chloride (DPI) of O_2_^·–^ to treat WT for 5 days after 1 DAG in 1/2 MS. In 250nM DPI- or 0.5mM PG-contained 1/2 MS, the root length was decreased dramatically comparable to that of *prl1* (Figure 2E and 2F). We stained PG-treated plant roots with DAB and NBT respectively and found similar distribution patterns of H_2_O_2_ and O_2_^·–^ to those of *prl1* over whole root (Figure 2D and 2G) implying ROS imbalance is enough to causing short root.

**Figure 2.**
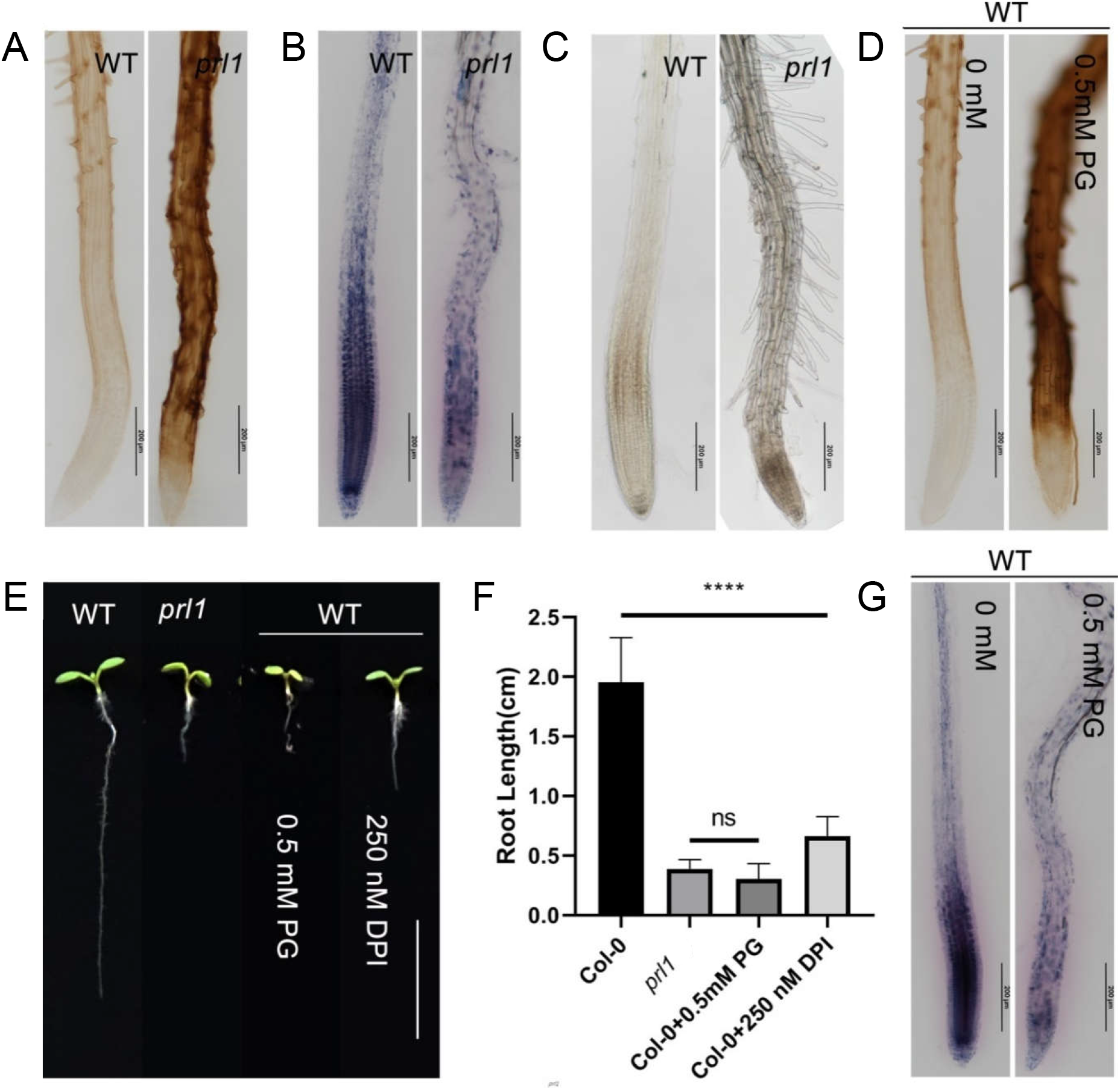
ROS imbalance in the root of *prl1*. (A) Roots stained with DAB marking distribution and concentration of H_2_O_2_ in WT and *prl1*. (B) Root stained with NBT indicating distribution and concentration of superoxide (O_2_^·–^). (C) Root hair along longitudinal axis of roots. (D) WT Roots stained with DAB grown for 5 days in none-PG-added and PG-added ½ MS. 5 days cultivation aimed for observing inhibitory effect of imbalance ROS on root development. (E) Roots of WT, *prl1*, PG-treated and DPI-treated WT. Scale Bar = 1cm. (F) Quantitative root length in WT, *prl1*, PG-treated and DPI-treated WT. (****p < 0.001, student’s test). (G) WT Roots stained with NBT for 5 days cultured in none-PG-added and PG-added ½ MS. Scale Bar in A, B, C, D and F is equal to 200 μm.

All together, we got that ROS was resettled to be imbalanced and its imbalance was enough to cause short root like *prl1*. **But, was ROS resettlement the necessary cause for short root of *prl1*?**

### ROS imbalance is undetermined the necessary cause for short root in prl1

The level of ROS in cells depends on the balance between ROS-generating and ROS-scavenging system (Eljebbawi et al., 2021; Mittler et al., 2004). To get knowledge about whether ROS imbalance is the reason for retarded primary root of the *prl1*, we utilized KI, a typical scavenger of H_2_O_2_, to remove hydrogen peroxide and resultantly restore cellular ROS balance (Tsukagoshi et al., 2010). We adopted two kind of criteria to evaluate the effectiveness of KI on root development, one was root length, and the other was DAB- and NBT-stained intensity and distribution patterns of H_2_O_2_ and O_2_ . After 1 day germinated in 1/2 MS, the WT and *prl1* seedlings were transferred into 1mM KI-containing 1/2 MS for further 5 days growth. As we saw, the KI-treated roots of WT but not of *prl1* were longer than their respective untreated counterparts indicating KI-treatment was effective for primary root elongation in WT but not *prl1* (Figure 3A and 3D). As for ROS, the DAB-staining intensity showed H_2_O_2_ concentration in KI-treated WT was reduced and its distribution shift upward; the NBT-staining marked O_2_^·–^ pattern also was changed with distribution to upward region (Figure 3B and 3C). These results were consistent to previous conclusion ROS regulates cell state transition from division to differentiation controlling root development (Tsukagoshi et al., 2010). Upon none effectiveness of KI on root length of *prl1*, we found H_2_O_2_ intensity was almost no difference between the *prl1* and the KI-treated *prl1* and the O_2_ ^· –^ distribution gradient was not distinguishably changed (Figure 3B and 3C). Taken together, these results indicated KI treatment could not effectively rebalance ROS in the root of *prl1*.

**Figure 3.**
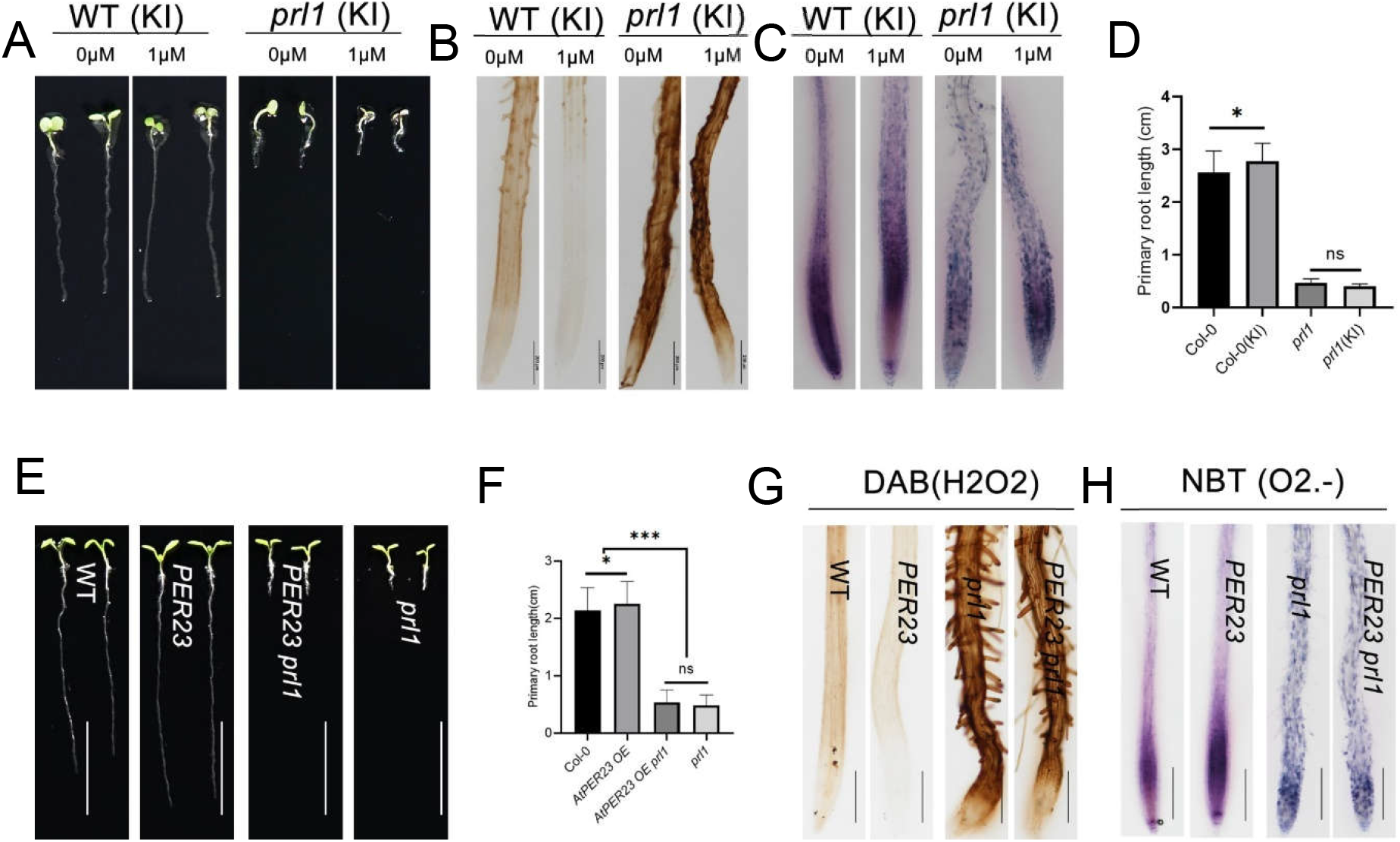
ROS imbalance is not determined the necessary cue for shortening root of *prl1*. (A) Representative roots of WT and *prl1* raised for 6 DAG in ½ MS or KI-containing ½ MS. DAB-stained roots of WT and *prl1* raised for 6 DAG in ½ MS or KI-containing ½ MS respectively. (C) NBT-stained roots of WT and *prl1* raised for 6 DAG in ½ MS or KI-containing ½ MS respectively. (D) Statistic analysis of root length of WT and *prl1* raised for 6 DAG in ½ MS or KI-containing ½ MS. (E) Representative 6-DAG roots of WT, *PER23OE* line, *PER23 prl1* and *prl1* raised in ½ MS. (F) Statistic analysis of root length of WT, *PER23*, *PER23 prl1*, and *prl1* raised for 6 DAG in ½ MS. (G) DAB-stained roots of WT, *PER23*, *PER23 prl1*, and *prl1* raised for 6 DAG in ½ MS. (H) NBT-stained roots of WT, *PER23*, *PER23 prl1*, and *prl1* raised for 6 DAG in ½ MS.

As chemical method could not effectively rebalance ROS in root of *prl1*, we turned to adopt genetic method to restore ROS balance. We performed RNA-seq with WT and *prl1* and analyzed transcriptional difference between WT and *prl1* about genes which would be correlated with ROS imbalance in *prl1*, then found multiple peroxidase-encoding genes (*PER*s) were down-regulated (Figure S2A). We utilized quantitative real-time PCR and RT-PCR to confirm these results (Figure S2B and 2C). According to collective knowledge in in TRAVADB (travadb.org), *PER23* was revealed to be wholly expressed in almost every type of root cell (Klepikova et al., 2015). To effectively clean out ROS (H_2_O_2_) and avoid disturbance to promoter activity of *PER23* by *prl1* mutation, we chose 35S promoter to drive *PER23* expression. Thereafter, we constructed homozygous double *PER23 prl1* mutants by cross. The expression levels of *PER23* in various *PER23OE* lines were confirmed by RT-qPCR to be higher than WT (Figure S2D). We chose *PER23OE* line with highest expression level among isolated lines to observe root length. We found this line took longer root than WT but the root length of *PER23 prl1* was like *prl1* (Figure 3E and 3F). As we expected, *PER23* overexpression line had lower H_2_O_2_ concentration than WT while the maximal concentration of O_2_^·–^ was distributed upward and its gradient kept similarity to WT (Figure 3G and 3F). The ROS patterns in the *PER23OE prl1* were kept like *prl1* (Figure 3G and 3H). These results indicated the constitutive overexpression of *PER23* could not effectively restored ROS balance in *prl1*.

Taken together, these results suggested that ROS imbalance is undetermined the necessary cause for short root of *prl1* implying other additional cues would generate ROS imbalance. Would the additional cues be the common cause of both short root and ROS imbalance in *prl1*?

### Stele cell death in *prl1* is independent of ATM/ATR-SOG1 module

Given that H_2_O_2_ is accumulated in the primary root of *prl1* and its accumulation is typically considered to be harmful to genomic stability by damaging DNA (Baxter et al., 2014; Gechev and Hille, 2005; Petrov et al., 2015), we surmised stele cell death in *prl1* would be the result of H_2_O_2_ accumulation-initiated DDR. We all knew that ATM/ATR-SOG1 module is a canonical signaling cascade mediated DDR in plant (Fulcher and Sablowski, 2009; Yoshiyama, 2016). We ordered T-DNA insertion mutants *atm* (SALK_040423c), *atr* (SALK_083543c) and *sog1* (SALK_039420) from ABRC, which were confirmed to be homozygous by specific primers (Table S3), and performed real time PCR to identify they all were null mutants (Figure S3). Subsequently, we crossed these mutants to *prl1* respectively and got the double homozygous *atm prl1*, *atr prl1* and *sog1 prl1* mutants. We found the 6-DAG roots of *atm*, *atr* and *sog1* were similar to WT (Figure 4A), but roots of the *atm prl1*, *atr prl1* and *sog1 prl1* were similar to those of *prl1* (Figure 4C) indicating genetically *PRL1* was epistatic to *ATM*, *ATR* and *SOG1* in regulation of root development. Using PI staining to mark dead cells, we found there was no stele cell death in *the atm*, *atr* and *sog1*, but in roots of the *atm prl1*, *atr prl1* and *sog1 prl1* there was still dead stele cells marked by symbol “※ (Figure 4B and 4D) suggesting that root stele cell death in *prl1* was independent of ATM/ATR-SOG1 module which further indicated ROS imbalance may not be the cause of root stele cell death.

**Figure 4.**
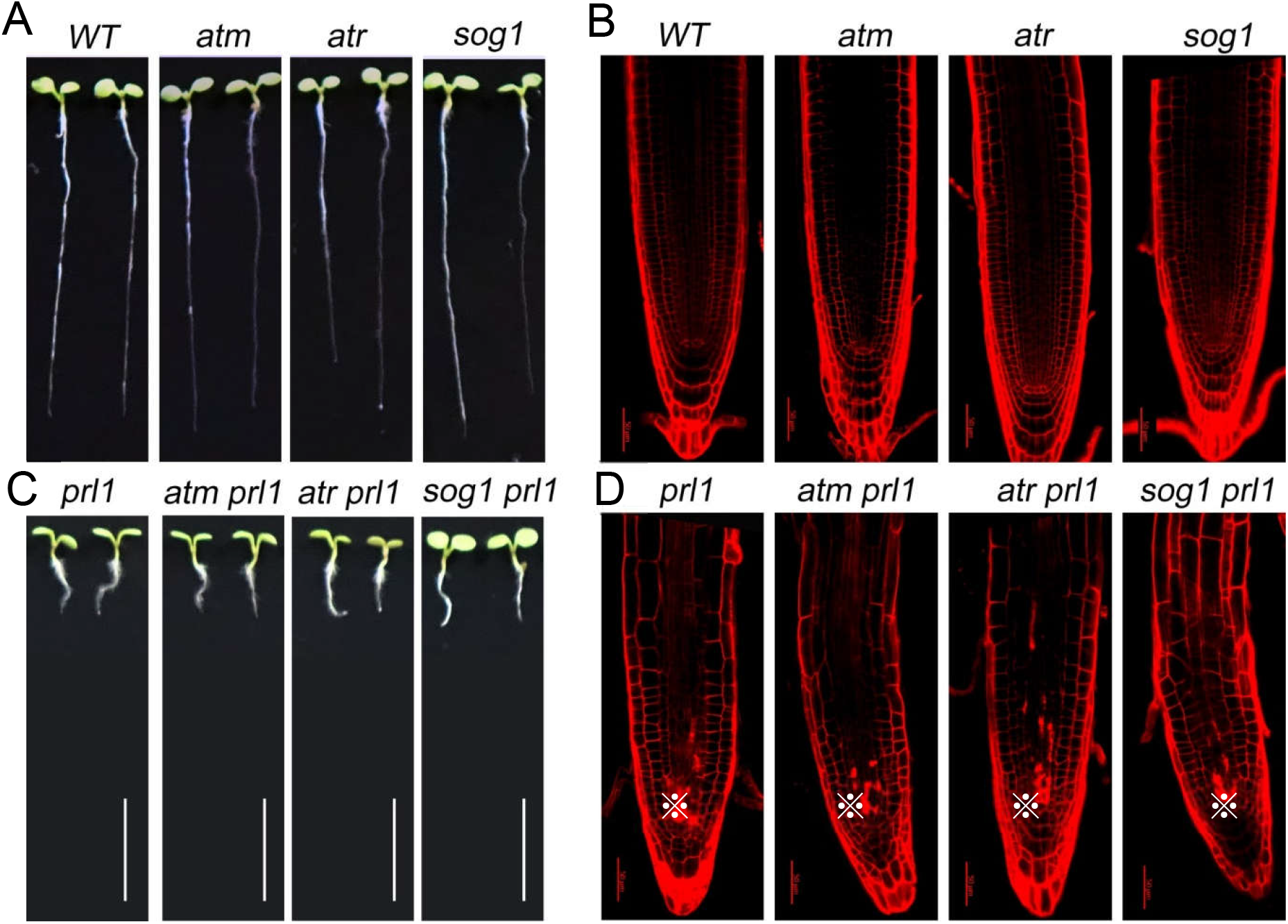
PRL1 is epistatic to ATM/ATR-SOG1 module on root growth and stele cell death. (A) Root length of WT, *atm*, *atr* and *sog1* in 6 DAG. (B) Root cell status of WT, *atm*, *atr* and *sog1* indicated by PI staining. (C)6-DAG root length of *prl1*, *atm prl1*, *atr prl1* and *sog1 prl1*. (D) Stele cell death in 6-DAG roots of *prl1*, *atm prl1*, *atr prl1* and *sog1prl1* indicated by PI staining. Scale bar in A and C equals to 1 cm, in B and D is 50 μm. “※” marks dead stele cell.

### Actin filament disassembly in *prl1* results in short root and ROS imbalance

Root length is a comprehensive developmental output of multiple signals. Our presented evidence is not enough for supporting ROS imbalance the causative factor of developmental defects of root in *prl1*. Fortunately, our work in deciphering mechanism for polar developmental defects of pavement cell in *prl1* enlightened us that cortical actin filament was depolymerized in pavement cells of *prl1*. Given actin as structural scaffold is the basic target of upstream signal and directly affects cell behaviors, we reasoned that MF de-polymerization would be the essential cause of developmental root defects in *prl1*. Utilizing confocal fluorescent microscope to observe cortical MF of epidermal cells in root elongation zone of constructed homozygous the *fABD2-GFP prl1* double mutant, we found that in the cortical region of WT cell was distributed with filamentous actin structure whereas in *prl1* mutant the actin configuration was failed to shape into filamentous structures (Figure 5A). Along the longitudinal axis of root, we checked and compared the cortical actin configurations in cells of the typical meristem zone, elongation zone and differential zone of *prl1* and WT. In WT, although actin filaments exhibit different patterns, cytoskeletal actin was observed to be filamentous structures in these different types of cells (Figure S4). In the three counterpart zones of *prl1* root cells, actin fails to form filamentous structures (Figure S4). These results suggested that actin filament structures were disrupted in *prl1*. To confirm that actin de-polymerization would be enough for short root, we employed actin polymerizing inhibitor LatB (1μM in 1/2 MS) to treat growing root. After 5 days treatment with LatB from 1 DAG, we got short-root seedlings like *prl1* (Figure 5B and 5E) and observed actin filaments to be completely de-polymerized in root cells of the LatB-treated seedlings (Figure 5A) suggesting de-polymerization of actin filament is enough for root growth retardation of *prl1*.

**Figure 5.**
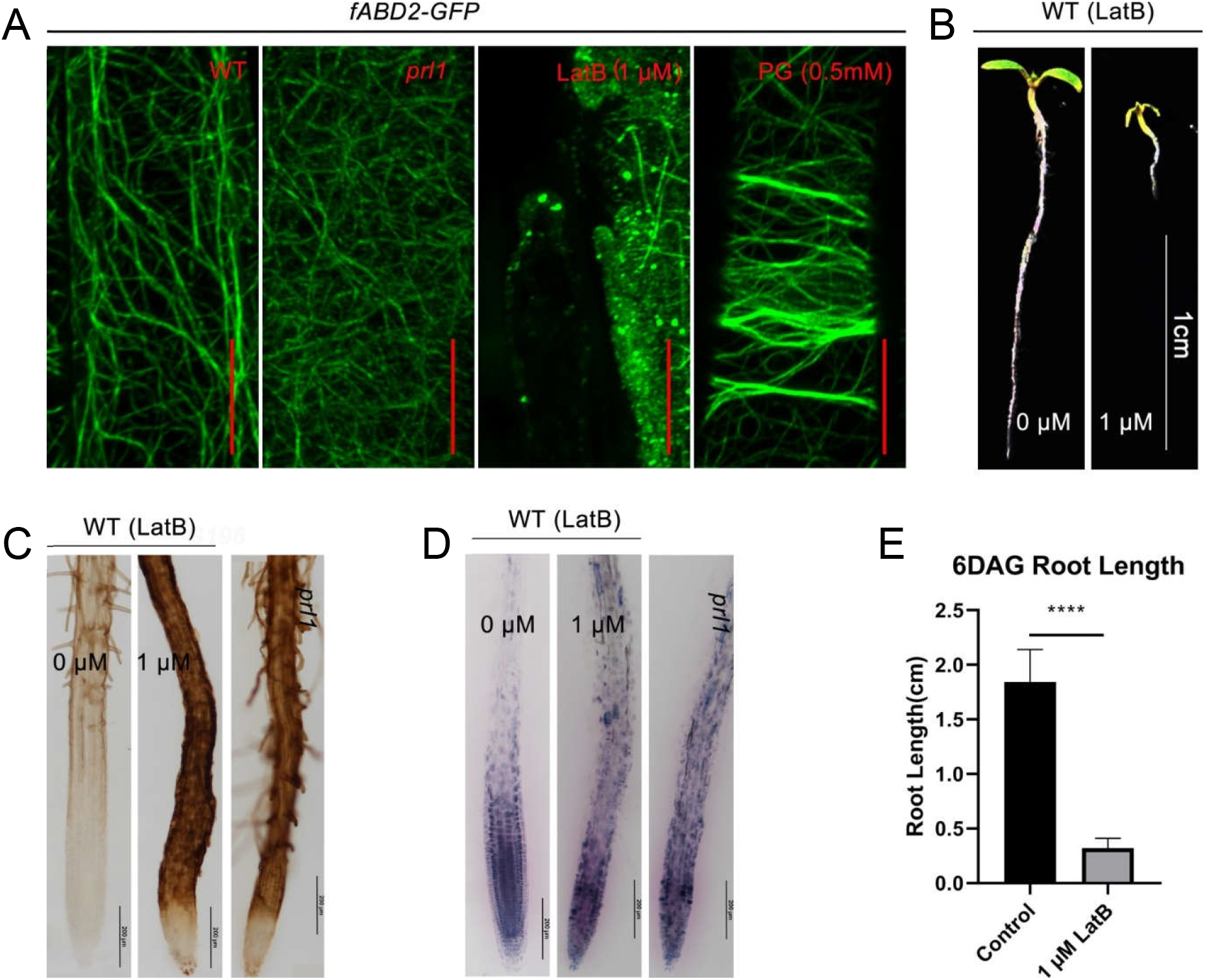
Actin disassembly is the fundamental cause for short root of *prl1* and results in ROS imbalance. (A) fABD2-GFP marked cortical actin structure in root EZ cells of WT, *prl1*, LatB(1μM)- and PG (0.5mM)-treated WT. In WT, filamentous actin structure is existing; in *prl1*, fluorescent intensity is faint and actin prefers to disassembly; in LatB-treated WT, actin filament is disappeared; in PG-treated WT, actin is formed into transversally arranged actin bundle. Scale bar = 10 μm. (B) Non and LatB-treated roots. 1-DAG seedlings were transferred into 0 μM or 1 μM LatB-containing ½ MS raised for 5 days. Scale bar = 1cM. (C) Roots of WT, LatB-treated WT and *prl1* stained with DAB. Scale bar equals to 200 μm. (D) Roots of WT, LatB-treated WT and *prl1* stained with NBT. Scale bar equals to 200 μm. (E) Quantification of root length of non and 1μM LatB-treated seedlings. Values are mean±SD of more than 30 roots at 6 DAG. Asterisk indicates significant difference (****p<0.0001, t test).

To clarify relationship between disassembled MF and ROS imbalance on defective development of the *prl1* root, we used DAB and NBT staining to detect levels of H_2_O_2_ and O_2_ ^· –^ and found that abnormal accumulation of hydrogen peroxide along primary root, concentration reduction and gradient disruption of superoxide in LatB-treated WT root like those in *prl1* (Figure 5C and 5D). Therefore, de-polymerization of MF would bring about both short root and ROS imbalance. To rule out the probability of ROS imbalance being the side effect of LatB-treatment and to confirm the conclusion of MF de-polymerization leading to ROS imbalance, we chose to observe effect of ROS imbalance on actin configuration. Again, 0.5mM PG were used to treat *fABD2-GFP* seedlings. In addition to observed short root (Figure 2E and 2F), we found cellular cortical MF were reorganized into transversally arranged thicker bundles in cells of elongation zone which were completely different to actin configuration in the *prl1* (Figure 5A), which indicated that ROS imbalance promoted formation of transverse actin bundle but not depolymerization of actin configuration and further suggested actin de-polymerization triggered ROS imbalance.

Taking together, actin de-polymerization is the leading cause of ROS imbalance and dominant to ROS imbalance being the cause of root development retardation of *prl1*. As cellular homeostasis is characterized by a baseline level of ROS (Mittler et al., 2022a), collectively it is concluded *PRL1* functions to maintaining cellular actin integrity and concomitant cellular homeostasis.

### ANAC085 specifically bridges MF de-polymerization signal to root stele cell death

Above evidence does not support stele cell death is the result of ROS accumulation-triggered and ATM/ATR-SOG1 signaling cascade-mediated DNA damage response. We turned to further analyze comparable RNA-SEQ data and found some *ANAC* genes exhibiting higher transcription levels in the *prl1* compared to WT. We performed RT-qPCR to quantify their expression levels (Fig. 6A). ANAC is a type of plant-specific transcription factor family including *SOG1*, whose members are always induced in response to stresses for plant resistance to stresses (Puranik et al., 2012). Referring to basic RNA-SEQ database in TRAVADB (travadb.org), we found *ANCA085*, normally expressed at very low levels in all tissues, was highly induced in *prl1*(Figure 6A and 6D). Then, we reasoned *ANAC085* would be specifically activated in response to *prl1* mutation-triggered ROS imbalance and mediated stele cell death. We took *ANAC044* as control, which has been also proved to be induced by stress (Takahashi et al., 2019). First, we got T-DNA insertion *anac044* (SALK_054551C) and *anac085* (SALK_208662C) mutants and analyzed genetic relationships between *ANAC044*, *ANAC085* and *PRL1* in root development via constructing *anac044 prl1* and *anac085 prl1* double homologous mutants. The root lengths of them were like that of *prl1* but significantly shorter than WT indicating *PRL1* was epistatic to *ANAC044* and *ANAC085* on root development (Figure S5). Second, we checked ROS levels in homologous *anac044 prl1* and *anac085 prl1* double mutants by DAB and NBT staining. The ROS distribution patterns and concentration in roots of the *anac044 prl1* and *anac085 prl1* were observed to be like in *prl1* (Figure 6B and 6C) suggesting ROS imbalance is independent of *ANAC044* and *ANAC085.* According to results of PI-stained root, we found there was not stele cell death in the root of *anac044* and the *anac085* like in WT roots. In roots of *anac085 prl1*, cell death was vanished or alleviated while it was still existing in the roots of *anac044 prl1* indicating *ANAC085* was required for *prl1* mutation-derived root stele cell death (Figure 6E). To dissect whether ROS imbalance was the cause of stele cell death and which was mediated by *ANAC085* in *prl1*, we utilized PG treatment of WT root for 5 days to mimic ROS status. WT seeds were germinated in 1/2 MS, 1 day after germinated, the seedlings were transferred to 0.5 mM PG-containing 1/2 MS, the treated roots at 6DAG were stained by PI. *ANAC085* was confirmed utilizing RT-qPCR to be up-regulated in PG treatment seedlings (Figure 6D). Out of our expectation, we observed no stele cell death was emerged in PG- and DPI-treated roots (Figure 6E). These results illustrated that ROS imbalance was not the cause to stele cell death in *prl1* and up-regulation of *ANAC085* alone was not enough to cause stele cell death. Then, we utilized actin polymerizing inhibitor LatB (1μM) to treat *anac085* and WT (*pWOX5::GFP* was used to conveniently observe position of QC) to learn role of actin de-polymerization in cell death, we found cell death emerged in the LatB-treated *pWOX5::GFP* but not the *anac085* mutant and the degree of cell death in *pWOX5::GFP* was increased with duration of LatB treatment (Figure 6E) suggesting actin depolymerization could be the cause of root stele cell death in *prl1*. Although we cannot make a distinction between ROS and actin de-polymerization about their role in induction of *ANAC085* expression, these results demonstrated that actin de-polymerization but not ROS imbalance was the essential cause of and in combination with *ANAC085* specifically caused stele cell death.

**Figure 6.**
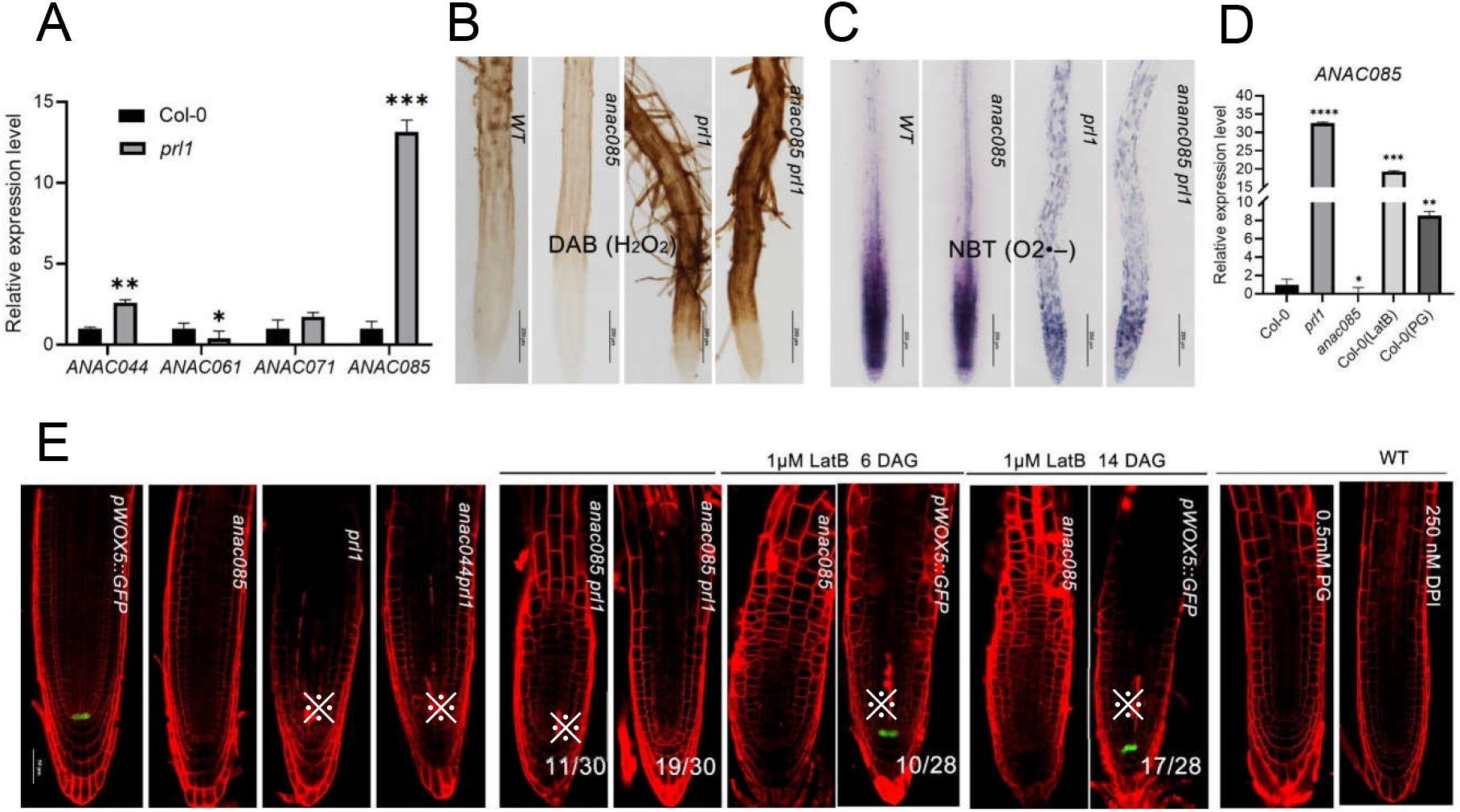
MF de-polymerization induces expression of and cooperates with *ANAC085* leading to stele cell death. (A) Relative expression level of ANAC TF members *ANAC044*, *ANAC061*, *ANAC071* and *ANAC085* in *WT* and *prl1*. In WT, their expression levels were normalized into “1”. (B)DAB-stained 6-DAG roots of WT, *anac085*, *prl1* and *anac085prl1*. Scale bar = 200 μm. (C)NBT-stained 6-DAG roots of WT, WT, *anac085*, *prl1* and *anac085prl1*. Scale bar = 200μm. (D)Relative expression level of *ANAC085* in WT, *prl1*, *anac085* and LatB-treated WT. The expression level of ANAC085 in WT was normalized into “1”. (E)PI staining indication of root stele cell living status in WT, *pWOX5::GFP*, *anac085*, *prl1*, *anac085prl1*, LatB-treated 6-DAG/14-DAG *anac085* and *pWOX5::GFP*, PG-treated and DPI-treated WT. *pWOX5::GFP* was used to indicate QC position. Scale bar = 50 μm. “※” marks dead stele cell.

In normal growth condition, single *anac085* and *anac044* has comparable phenotype to WT respectively, but in the genetic background of *prl1*, defects in *ANAC085* and *ANAC044* has even more serious seedlings growth inhibition at 11 DAG (Figure S5) than *prl1* supporting above conclusion *prl1* mutation triggered stresses and reaffirmed the common roles of *ANAC085* and *ANAC044* in making trade-off between stress response and plant growth.

### Differentially expressed genes (DEGs) and alternative splicing (AS) are mutually independent events in response to actin depolymerization in *prl1*

PRL1 has widespread been confirmed to repress stress-related gene transcription (Jia et al., 2017; Nemeth et al., 1998; Salchert et al., 1998). To confirm relationship between *prl1* mutation-caused actin de-polymerization and transcriptional alteration, we performed comparable RNA-seq analysis with *prl1*, LatB-treated WT, and WT in three biological replicates. The significant thresholds for differentially expressed genes (DEGs) between each experimental group and WT were settled with |log2FoldChange|≥1 and padj<0.05. We identified 1932 DEGs with 714 up-regulated (hyper-DEGs) and 1218 down-regulated (hypo-DEGs) in *prl1* (Fig. S7 A, Data Set S1); 3027 DEGs with 1596 hyper-DEGs and 1431 hypo-DEGs in the LatB-treated WT (Fig. S7 B, Data Set S2). we found there were 800 common DEGs between *prl1* and LatB-treated WT and the overlap of the DEGs was significant (Fisher’s exact test, P=0) (Fig.7A) indicating *prl1* and actin depolymerization regulated the same group of genes and shared similar actin depolymerization-related signaling pathway. To analyze their functional correlations, we performed GO enrichment on respective DEGs of them. The DEGs in two independent experiments were enriched in stimulus responses, including response to toxic substance, response to chemical and response to stress indicating *PRL1* mutation and actin depolymerization share downstream signaling pathways related to stresses (Fig. 7D and 7E), further indicating PRL1 performs biological functions through maintains actin integrity and cellular homeostasis. In *prl1* and LatB-treated WT, DEGs with biological function related to oxidative stress is remarkable (Fig. 7D and 7E). ROS imbalance was the common result of *prl1* and actin depolymerization, we analyzed DEGs in PG-treated WT compared to WT and identified DEGs with 795 hyper-DEGs and 1724 hypo-DEGs (Fig. S7 C, Data Set S3). Then, we checked their intersections and got 549 DEGs were overlapped and the overlap level was significant (SuperExactTest, P=0) (Fig. 7C) indicating quite a few of downstream stress-related transcriptional alteration was directly responsive to ROS imbalance (Fig. 7F). This result demonstrated DEGs were not directly related to PRL1-regulated transcriptional machine but the responses to actin depolymerization- or ROS imbalance-related stresses. In addition to transcription regulation, *PRL1* and it-localized MAC/Prp19 complex had been revealed in function of regulating alternative splicing (AS) by regulation of activation of spliceosome complex (Chanarat and Sträßer, 2013), but the underlying mechanism is unknown. Given relationship between stress and AS defect as AS is always the adapted strategies for plants to respond to stresses (Laloum et al., 2018; Ling et al., 2021), we proposed AS defect in *prl1* was the event of actin depolymerization-originated stress. Seeing that intron retention (RI) is a major kind of alternative splicing defect in Arabidopsis (Ner-Gaon et al., 2004), we analyzed RI in both *prl1* and LatB-treated WT compared to WT and found that 664 RI was in *prl1* and 401 RI in LatB-treated WT. The number of overlapped RI was 187 and the overlap level was statistically significant (Fisher’s exact test, P=0) (Fig. 7B) indicating RI defects in *prl1* came from actin depolymerization. To reveal biological functions of the overlapped RI defect genes, we performed GO enrichment to them and found the genes enriched were components of the spliceosome complex (Fig. 7G) including splicing regulator SR30 (AT1G09140), SR33 (AT1G55310), SUA (AT3G54230) and so on (Data Set S4), indicating *prl1* and actin depolymerization-shared AS defect (RI) genes were alternative splicing regulators. Increasing evidence indicates that biotic and abiotic stresses could change the alternative splicing of splicing regulators (Reddy, 2007), hence PRL1 mutation-initiated stresses would be reasonable the cause of alternative splicing defect. Thereafter, DEGs and AS defects in *prl1* are also the result of actin depolymerization-initiated stresses. To reveal whether there is any correlation between splice variant and gene expression in *prl1* and LatB-treated WT, we analyzed the expression distribution of intron-retained genes and found them randomly distributed in increased, reduced and unchanged regions indicating there was no correlation between intron retention defects in *prl1* and LatB-treated WT (Fig. 7H and 7G) which is consistent to Jiàs result (Jia et al., 2017). Come together, DEGs and AS defects (RI) are the parallel responses to actin depolymerization in *prl1*.

**Figure 7.**
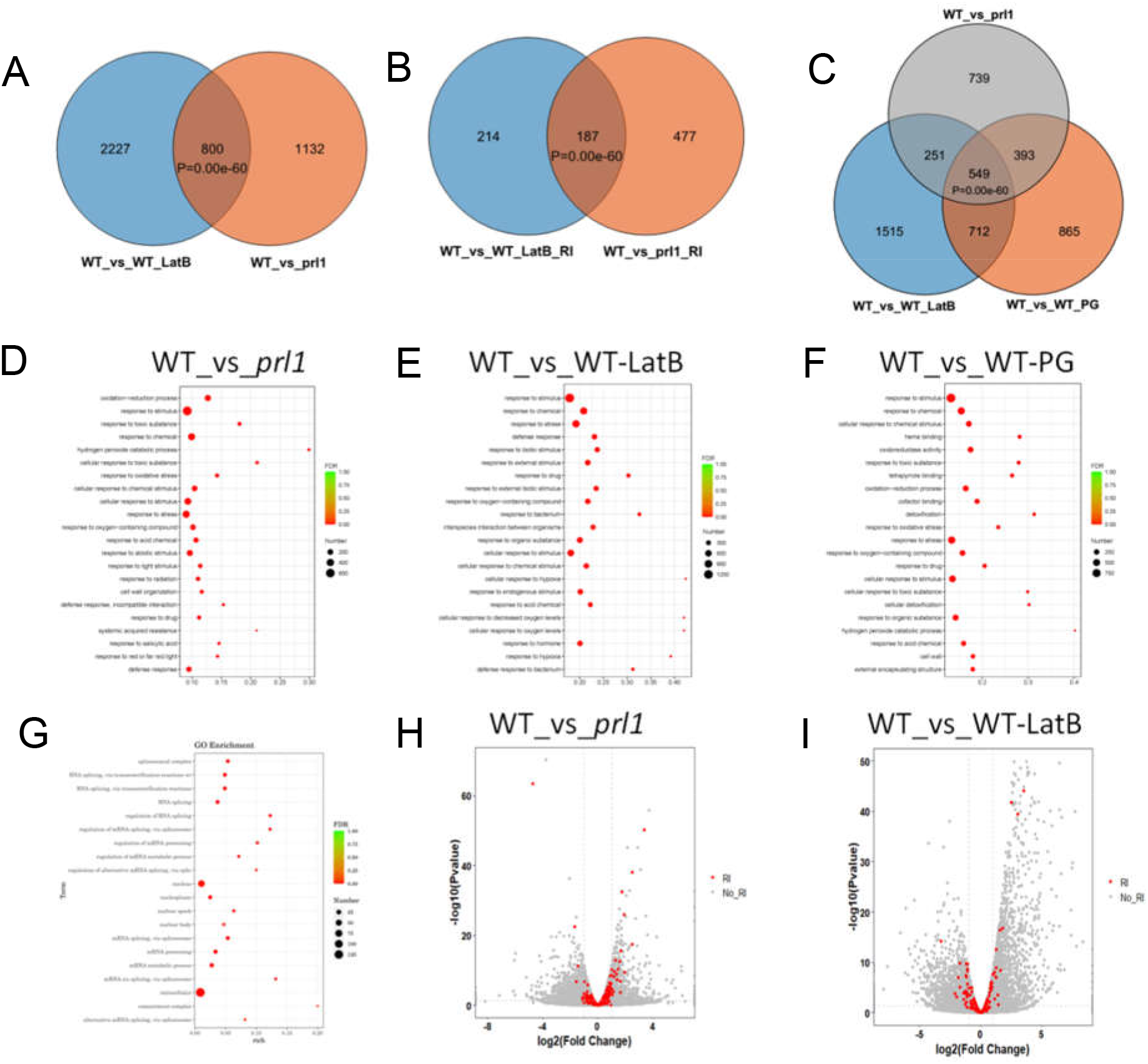
PRL1 negatively regulates actin de-polymerization and resultantly stress-correlated differential gene expression. (A) A Venn diagram showing the degree of overlap among differentially expressed genes (DEGs) in *prl1* and LatB-treated WT (Fisher’s Exact test, P=0). (B) A Venn diagram showing the degree of overlap among genes with retained intron (RI) in *prl1* and LatB-treated WT (Fisher’s Exact test, P=0). (C) A Venn diagram showing the degree of overlap among differentially expressed genes (DEGs) in *prl1*, LatB-treated WT and PG-treated WT (SuperExactTest, P=0). (D) GO enrichment analysis of DEGs in *prl1* showing they are stress-related. (E) GO enrichment analysis of DEGs in Lat-treated WT showing they are stress-related. (F) GO enrichment analysis of DEGs in PG-treated WT showing they are stress-related. (G) GO enrichment analysis of overlapped intron retention genes in *prl1* and LatB-treated WT showing they are RNA splicing-related. (H) Volcano plots showing fold changes of gene expression levels in *prl1* compared with WT, red dots indicating intron retention genes, grey dots indicating DEGs. Significant thresholds for fold change with |log2FoldChange|≥1 and padj≤0.05 are shown in the plot as gray dashed lines. (I) Volcano plots showing fold changes of intron retention genes in LatB-treated WT compared with WT. Significant thresholds for fold change with |log2FoldChange|≥1 and padj≤0.05 are shown in the plot as gray dashed lines.

## Discussion

### Cellular structure (actin de-polymerization) is dominant to signal (ROS imbalance) on retarded root development

Actin cytoskeleton is experimentally evidenced and well accepted to be the basic determinant of cell behaviors such as cell division, cell expansion, cell morphogenesis and vesicle transport (Hussey et al., 2006; Kadzik et al., 2020). Root length is the combinatory result of cell proliferation and directional cell elongation (Beemster and Baskin, 1998; Takatsuka et al., 2018; Vaughn et al., 2011b). During mitosis of plant cells, actin is dynamic changing from one form to other to perform temporally specific cell division functions (Vaughn et al., 2011b). When cells reach the elongation zone, they will expand directionally in parallel with longitudinal axis of primary root (Dolan and Davies, 2004; Takatsuka et al., 2018). Both exogenous and endogenous cues have effects on root development. The actin cytoskeleton is typically considered the target of upstream signals which modulate ABPs (Actin binding proteins) for actin configuration turnover (Huang et al., 2011). So, actin de-polymerization is the final determinant of short primary root in the *prl1* mutant.

ROS imbalance is produced in response to various internal or external stress signals, which in turn affects series of processes of cell including cell physiology, cell signaling, membrane properties, cell wall relaxation and final cell death (Mittler et al., 2022b). Therefore, ROS imbalance can indirectly suppress root growth. Either ROS imbalance or actin de-polymerization could be the enough cause to short root of *prl1* alone. In our work, we gradually deduced some given factor preceded the ROS imbalance leading to stele cell death of *prl1*. Further, we found actin was depolymerized and confirmed its de-polymerization to be the fundamental cause of both ROS imbalance and short root of *prl1*. Our work presented ROS imbalance could made feedback on actin reorganization but failed in rescuing its depolymerized status in *prl1*. In view of effect of actin structures on root growth inhibition, our work indicated that ROS imbalance could reorganize configuration and spatial distribution of actin cytoskeleton which is different to actin depolymerization in *prl1*. In this work, the key for making forward step about deciphering mechanism for *PRL1* in root development is the finding of de-polymerization of cortical actin skeleton and discriminating between actin de- polymerization and ROS imbalance the essential cause to root developmental defect in *prl1*. Recently, a handful of evidence also suggested the actin reconfiguration triggered dangerous/stress signal which concomitantly produce ROS imbalance (Franklin-Tong and Gourlay, 2008; Gourlay and Ayscough, 2005; Thomas and Franklin-Tong, 2004).

### PRL1 inhibits stele cell death through maintaining dynamic actin homeostasis and cellular homeostasis

In both plant and animal cells, various cellular skeleton structures are important for multiple specific cellular functions such as directional outgrowth, vesicle transport, and cell morphogenesis. Further, it is realized that actin dynamic homeostasis (actin cytoskeleton is free of any bias to polymerization or de-polymerization) is more important than a single type of actin configuration for its function. In response to exogenous and endogenous signals, actin turnover is kept constant between G-actin and F-actin (Kadzik et al., 2020). About elaborate control of dynamic actin equilibrium, there are also many questions to be answered: how actin configurations are sensed during their being altered? What kinds of signals are produced from the instantaneous or the lasting alterations of actin configuration itself? Increasing data indicated actin cytoskeleton reorganization was recognized as dangerous signal (Desouza et al., 2012; Franklin-Tong and Gourlay, 2008; Gourlay and Ayscough, 2005; Leontovyčová et al., 2020; Thomas and Franklin-Tong, 2004). In self-incompatible plants, S-protein induces actin de-polymerization which in turn triggers caspase-involved death of self-incompatible pollen, DNA fragmentation and ROS accumulation which perform multiple-tier inhibition of pollination (Thomas and Franklin-Tong, 2004; Thomas et al., 2006). In mammalian and yeast cells, actin de-polymerization caused cell death and cell aging which is universally accompanied with but independent of ROS accumulation (Brown, 2012; Desouza et al., 2012; Gourlay and Ayscough, 2005). In consistent with previous work, we testified *prl1* mutation led to de-polymerization of actin cytoskeleton which in turn brought about cell death and irremovable ROS accumulation. Cellular homeostasis is always characterized by the level of ROS (Mittler et al., 2022b), in our work we found ROS imbalance in *prl1* mutant suggesting its cell out of homeostasis. Given ROS imbalance could lead to cell death in many cases, it is hard to distinguish its causative roles from actin de-polymerization in cell death, fortunately in our work we found *ANAC085* could specifically mediate signal from and functions in combination with the actin de-polymerization rather than ROS accumulation to cell death enforcing actin de-polymerization is the root cause of cell death. As *ANAC085* could be activated among *prl1*, LatB-treated WT and PG-treated WT, ROS imbalance could not be completely excluded to be involved in cell death in *prl1*.

### What is the Role of PRL1 in regulation of gene transcription?

In Previous works, by analyzing mutants, as transcriptional levels of many genes were found to be altered including mRNA, microRNA and siRNA, PRL1 and its orthologs *Prp46p* and *PLRG1* were deduced to be involved in regulating gene transcription (Chanarat and Sträßer, 2013; Hogg et al., 2010; Koncz et al., 2012). Using immune-affinity method, physical interaction between molecular components of Prp19/NTC and components of spliceosome complex or RNA biogenesis and processing machinery provided indirect evidence suggestion that PRL1 and its orthologs are involved in regulating gene transcription indirectly through interacting with component of transcriptional machinery (Koncz et al., 2012; Monaghan et al., 2009). As differential expression of genes usually drives dramatic impact on physiologies of organism, defects in innate immunity and development of the *prl1* were in haste deduced to be the effect of genes transcription alterations (Jia et al., 2017; Li et al., 2018; Monaghan et al., 2009; Palma et al., 2007). However, in our work we presented new discoveries and clarified that the depolymerization of actin cytoskeleton was the fundamentally more direct cause of developmental defect of primary root and reasonably deduced to be the cause of defects in innate immunity of *prl1* as actin de-polymerization was popularly accepted to be the cause of failure in resistance to pathogens invasion (Day et al., 2011; Janda et al., 2014; Jelenska et al., 2014; Leontovyčová et al., 2020; Porter et al., 2012; Tian et al., 2009). In addition, Jia and colleagues defined alternative splicing and differential expression of genes were mutually independent in three MAC subunit mutants including *prl1 prl2* mutant (Jia et al., 2017). Moreover, one intriguing question is why the de-repressed genes in *prl1* are determined to be biased to stress-correlated if PRL1 is involved in regulating transcriptional machinery to govern gene expression? Our biological GO terms indicated the DEGs were attributed to actin depolymerization-initiated stress/dangerous signal. So, in combination our results with previous works, we present evidence support the conclusion that transcriptional genes alterations in the *prl1* are responses to actin depolymerization-derived stress/dangerous signal.

Here, according to our present evidence we propose functional model of *PRL1*. Mutation in *PRL1* leads to cell cortical actin depolymerization, which results in growth and development defect of *prl1* with short primary root; concomitantly, actin depolymerization is recognized by cells to be stress/dangerous signal which initiates hierarchical events such as ROS imbalance, DEGs, AS defect and cell death which would make feedback on dynamic actin cycle and plant development et al., hence, our result presents a new scene of the function of *PRL1* in maintaining actin filament integrity and concomitant cellular homeostasis (Fig. 8).

**Figure 8.**
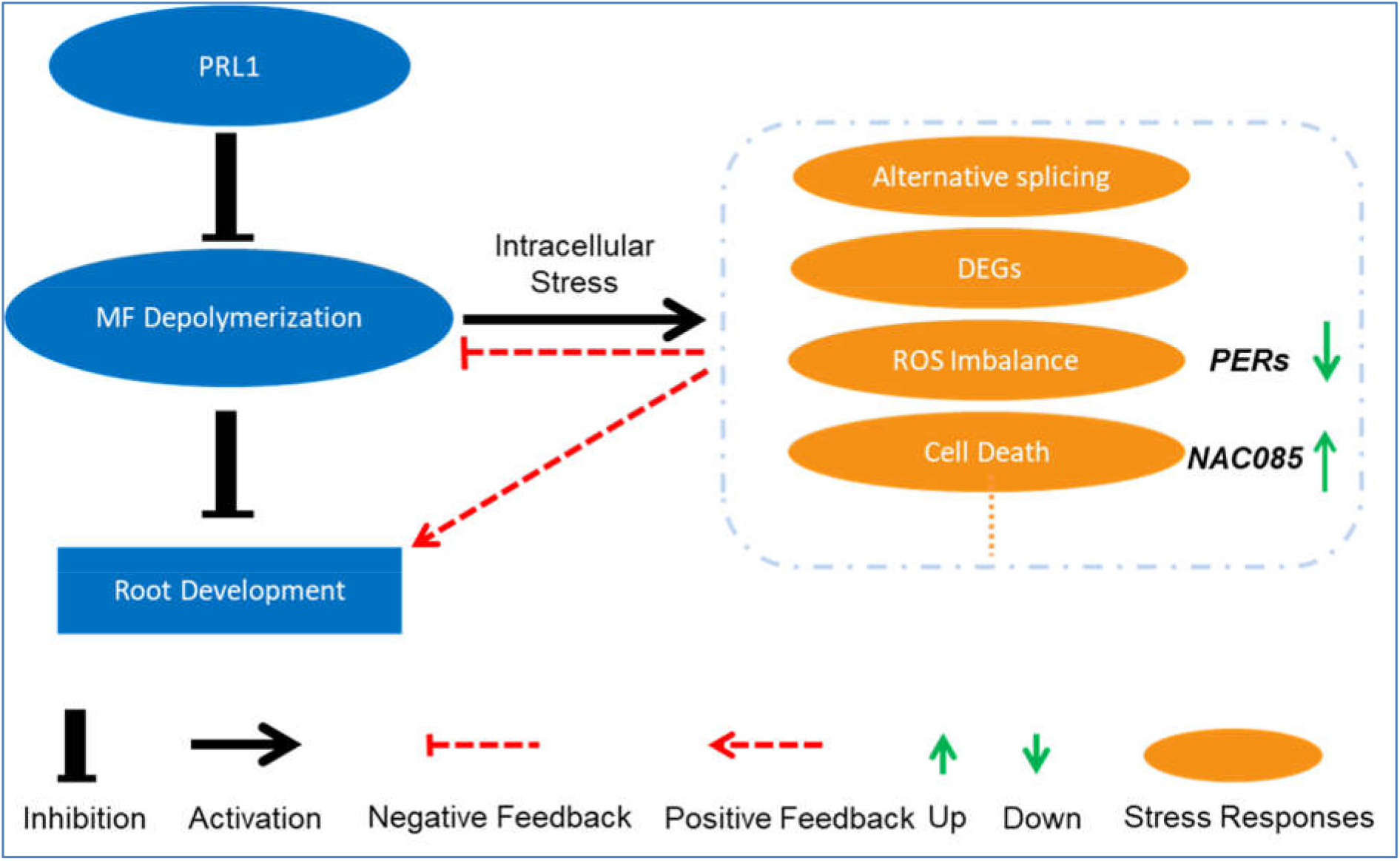
PRL1 maintains dynamic MF homeostasis and concomitantly cellular homeostasis. Actin depolymerization results in growth inhibition and development defect of *prl1* with short primary root; concomitantly, actin depolymerization is recognized by cells to be stress/dangerous-related signal which initiates hierarchical events such as ROS imbalance, DEGs, AS defect and cell death which would make feedback on dynamic actin cycle and plant development et al., hence, our result presents a new scene of the function of *PRL1* maintaining actin filament integrity and concomitantly cellular homeostasis.

## Material and Methods

### Plant materials and growth conditions

All the *Arabidopsis thaliana* plants used in this study are in the background of Columbia (Col-0) ecotype. The used *prl1* allelic mutants were uniformly marked *prl1* in the paper for convenience including a single nucleotide substitution mutant (*prl1-9*, transition of the 1042^th^ base C to T in the 13^th^ exon of *PRL1*) and a T-DNA insertional line Salk_039427; all the T-DNA insertion mutants were obtained from Arabidopsis Biological Resource Center (ABRC) including Salk_039427(*prl1-11*), Salk_054551(*anac044*), Salk_208662(*anac085*), Salk_039420(*sog1*), Salk_040423(*atm*), and Salk_083543(*atr*) were confirmed using PCR with primers listed in table S1. The *QC25*: *GUS (Sabatini et al., 1999)*, *pWOX5*:: GFP (Haecker et al., 2004) and *CycB1;1*: *GUS* (Colón-Carmona et al., 1999) seeds were kindly provided by Dr. CY, Li (IGDB, CAS, China). The seeds of actin marker line *pfABD2*: *GFP* was kindly provided by Dr. DS, Lin (HIST, FAFU, China). All used double mutants, *pWOX5*::*GFP prl1*, *pfABD2*::*GFP prl1*, p*35S*::*PER23*, *p35S*::*PER23 prl1 anac085 prl1*, *anac044 prl1*, *atm prl1*, *atr prl1*, *sog1 prl1* and *cycB1;1*:*GUS prl1* were got by cross and checked using PCR primers listed in table S1 and S3. Seedlings were germinated on ½ MS agar plates added with 1% sucrose in the growth incubator at 22℃ with 16hr light and 8hr dark.

### Construction and screen of *p35S::PER23* over-expression lines

For construction of the *p35S::PER23-GFP* over-expression lines, the coding sequence of *PER23* (AT2G38390) was cloned by PCR primers (Table S2) and constructed into pDONR207 by BP reaction, sequenced to make sure insertional fragment to be match to referred sequence and cloned it into pGWB605 by LR reaction, then the positive *p35S::PER23-GFP* was transferred into Agrobacterium tumefaciens GV3101 using the routine floral dip method (Zhang et al., 2006). The T1 seeds were broadcast in the soil, when germinated for 2 days, spayed with diluent Basta several times, the survival seedlings were transplanted to new pot. Seeds of transgenic lines was yield individually and the homozygous transgenic lines were screened in T2 and T3 generations. The selected homozygous transformants were used for followed experiments.

### Chemical pretreatment of seedlings

To evaluate the relationship between ROS and actin depolymerization and their effects on root development, the germinated seeds in ½ MS medium for only 1 day were transferred to PG- (0.5 mM), DPI- (0.25 µM), KI- (1 mM) and LatB-containing (1 µM) ½ MS medium separately and raised 5 days further. The 6 DAG seedlings were used for followed root length evaluation, ROS staining, actin configuration imaging and RNA-seq et al.

### Phenotypic analysis of root length

For root length measurements, the 6 DAG roots of WT or homologous mutant lines were photographed with while growing on agar (Willemsen et al., 1998). The root length data were collected using the ImageJ software and analyzed with statistic software GraphPad Prism 8 with 3 biological repeats.

### Root zones and cortical cell number analysis

Root tips of seedlings were photographed with DIC optic on an Eclipse Nikon Ni-U Upright Microscope. The number (root meristem cell number is defined as the number of cells in the cortex file extending from the QC to the transition zone) and length of epidermal cells were analyzed using Image J (Willemsen et al., 1998).

### Root PI staining and QC analysis (cell death observation)

Roots were stained with 10µg/mL PI (Sigma) for 5-10 min before imaging (Fulcher and Sablowski, 2009). After staining, the roots of *pWOX5::GFP* and *pWOX5::GFP prl1* were mounted on glass slides in distilled water. Confocal imaging was performed with Zeiss LSM880. GFP and PI were excited using 488 nm and 561 nm laser light individually. The emitted fluorescent light was collected between 500 and 550 nm (GFP) and 600 and 656 nm (PI). Z stacks were obtained by imaging 0.5-µm sections, which were averaged two times. Images were processed using Zeiss confocal software and Adobe Photoshop CC 2018.

### Beta-Glucuronidase (GUS) staining (Root cell division activity analysis (CycB1;1: GUS gus staining, QC25: GUS staining))

Seedlings with 6 DAG including *CycB1;1:GUS*, *CycB1;1:GUS prl1*, *QC25:GUS* and *QC25:GUS prl1* grown on ½ MS media in light were used for GUS staining. All roots were immersed in GUS-staining solutions (0.1M NaPO_4_ Ph7.0, 10mM EDTA, 0.1% (V/V) Triton X-100, 1 mM K_3_Fe (CN)_6_, 2 mM X-Gluc) for 2h to overnight at 37 ℃’in incubator, then washed 3 times using 75% ethanol solution until root tissue clear (Aida et al., 2004; Bieleszová et al., 2019). Then the roots were observed with a Nikon Ni-U Upright Microscope and the pictures were taken with Nickon Ni-U Upright Microscope equipped with a Digital Slight 10 camera.

### Starch granules (Lugol) staining

The root of the 6 DAG seedlings were submerged in Lugol solution (Sigma) for 5min, rinsed with distilled water, cleared with clearing buffer (chloral hydrate:glycerol:water in 8:3:1 ratio) (Aida et al., 2004), then observed and photographed using a DIC optic on a Nikon Ni-U upright Microscope equipped with a Nikon Digital Slight 10 Camera.

### NBT staining

Whole seedlings were used 6 day after germination. Roots were stained for 10-15 min in a solution of 2 mM nitroblue tetrazolium (NBT) in 20 µM phosphate buffer pH 6.1. The reaction was stopped by transferring the seedlings in distilled water (Dunand et al., 2007; Jia et al., 2017). The roots were observed with an Eclipse Nikon Ni-U Upright Microscope. Pictures were taken with Nikon Ni-U Upright Microscope equipped with a Digital Slight 10 camera. Settings were identical for all the pictures in an experiment. Each experiment was repeated at least 3 times with similar results.

### DAB staining

Whole seedlings were used 6 day after germination. Placed whole seedlings in freshly prepared DAB solution (Dissolve 10 mg 3,3’-diaminobenzidine (DAB) in 10ml distilled water, adjust the PH to 3.8 using HCl.) for 4 h in the dark at room temperature, remove the staining buffer and wash twice with ddH_2_O (Jia et al., 2017), observe and image with the H2O2 production and distribution in the roots with Nikon Ni-U Upright Microscope equipped with a Digital Slight 10 camera. Settings were identical for all the pictures in an experiment. Each experiment was repeated at 3 times.

### Live imaging of the cortical actin configuration using Confocal Microscopy

To visualize the cortical actin configuration, laser-scanning confocal microscopy LSM 880 with Airyscan to image the epidermal cells in different zones of the primary roots of *fABD2- GFP*, PG- and LatB-treated *fABD2-GFP* and *fABD2-GFP prl1* individually with identical parameters. Roots grown to 6 DAG were mounted on the slide, added distilled water and covered with coverslip and observed under Zeiss LSM 880. Took Z stacks of optical sections and Z-series maximum intensity projections of GFP fluorescence was generated to evaluate actin configurations.

### Real-time RT qPCR

RNA was extracted using TRIzol Reagent (Invirtrogen) from the seedlings of WT, *prl1*, *anac085*, *PER23* over-expressing lines, the LatB- and PG-treated WT respectively. First-strand cDNA was synthesized using the SuperScript III cDNA Synthesis kit (Invitrogen). RT qPCR was performed using 2×Hieff qPCR SYBR Green Master Mix (Yeasen, Shanghai, China) and PCR reactions and fluorescence detection were performed in a Mastercycler ep realplex (Eppendorf, New York, NY). ACTIN 2 (ACT2) was used as the internal control. Three technical replicates of the RT-qPCR were performed using three biological replicates. Primers used for qPCR are listed in Table S3.

### RNA-SEQ analysis

RNA-seq analysis was carried out at the Personal Biotechnology Co., Ltd (Shanghai, China). The total RNAs were extracted using the RNeasy Plant Mini Kit (Qiagen). The RNA-seq libraries were constructed using the TruSeq RNA Sample Prep Kit (Illumina). The mRNA was purified using oligo(dT) magnetic beads and then fragmented to about 300bp. The first-strand cDNA was synthesized with random hexamer primers by reverse transcription. The double-stranded cDNA was synthesized on template of single-fragment cDNA. The fragments with adaptors were amplified by adaptor-specific primers to enrich the fragment of library. The DNA library quality was tested by an Agilent 2100 Bioanalyzer, then check the total and effective concentration of DNA library. The libraries were applied to the Illumina Next-generation Sequencing platform for 150-nt Paired-end sequencing. Three biological replicates were performed separately. For each sample, the raw reads number was over 38 million. After filtration, the clean reads occupied over 92% of sequenced reads. The filtration steps included:

(1) remove reads containing only the adaptor sequences; (2) remove reads with average quantity lower than molecular quantity 20 (Q20). About 96% of the useful reads could be uniquely mapped to the reference of Arabidopsis thaliana TAIR10 using HISAT2 (http://ccb.jhu.edu/software/hisat2/index.shtml). At least 99.3% of the clean reads could be mapped to the exons, and about 99.0% clean reads could be mapped to genes. Gene annotation was referred to databases of Ensembl (http://www.ensembl.org/). Gene expression levels were normalized based on FPKM (fragments per kilobase of exon model per million mapped fragments). DESeq was used for analyzing the significantly differentially expressed genes (DEGs; |log2 FoldChange|>1, P-value <0.5). Analyzed the numbers of common and specific DEGs, constructed venn diagram of DEGs, used Fisher’s Exact Test to evaluate overlapping significance of DEGs between mutually experimental groups and SuperExactTest among 3 experimental groups about their functioning intersection. The program topGO was used for GO enrichment analysis, found significantly enriched GO terms. The software rMATs (http://rnaseq-mats.sourceforge.net/index.html) was used for analyzing Alternative Splicing (AS) events in the experimental samples. The quantification of AS events was conducted with rMATS statistic model. Δф (exon inclusion level) was used to be the standard confirming production of AS between two experimental samples (Δф>5% and FDR<= 1%). We also used Fisher’s Exact Test to evaluate overlapping significance of genes with RI (Retained Intron) events between *prl1* and LatB-WT experimental groups (Jia et al., 2017; Peng et al., 2022).

## Author contributions

X.G and Z.Y. conceived and designed the study. X.G., C.W. and X.W. performed the experiments and made data analysis. X.G. wrote the manuscript. All authors read and approved the final version of the manuscript.

## Supporting information

Supplemental Data Set 1

Supplemental Data Set 2

Supplemental Data Set 3

Supplemental Data Set 4

## Acknowledgments

We thank Wenwei Lin, Xu Chen, Tongda, Xu for comments and discussions.

## SUPPLEMENTARY MATERIALS

### Figure S1-S6

**Figure S1.**
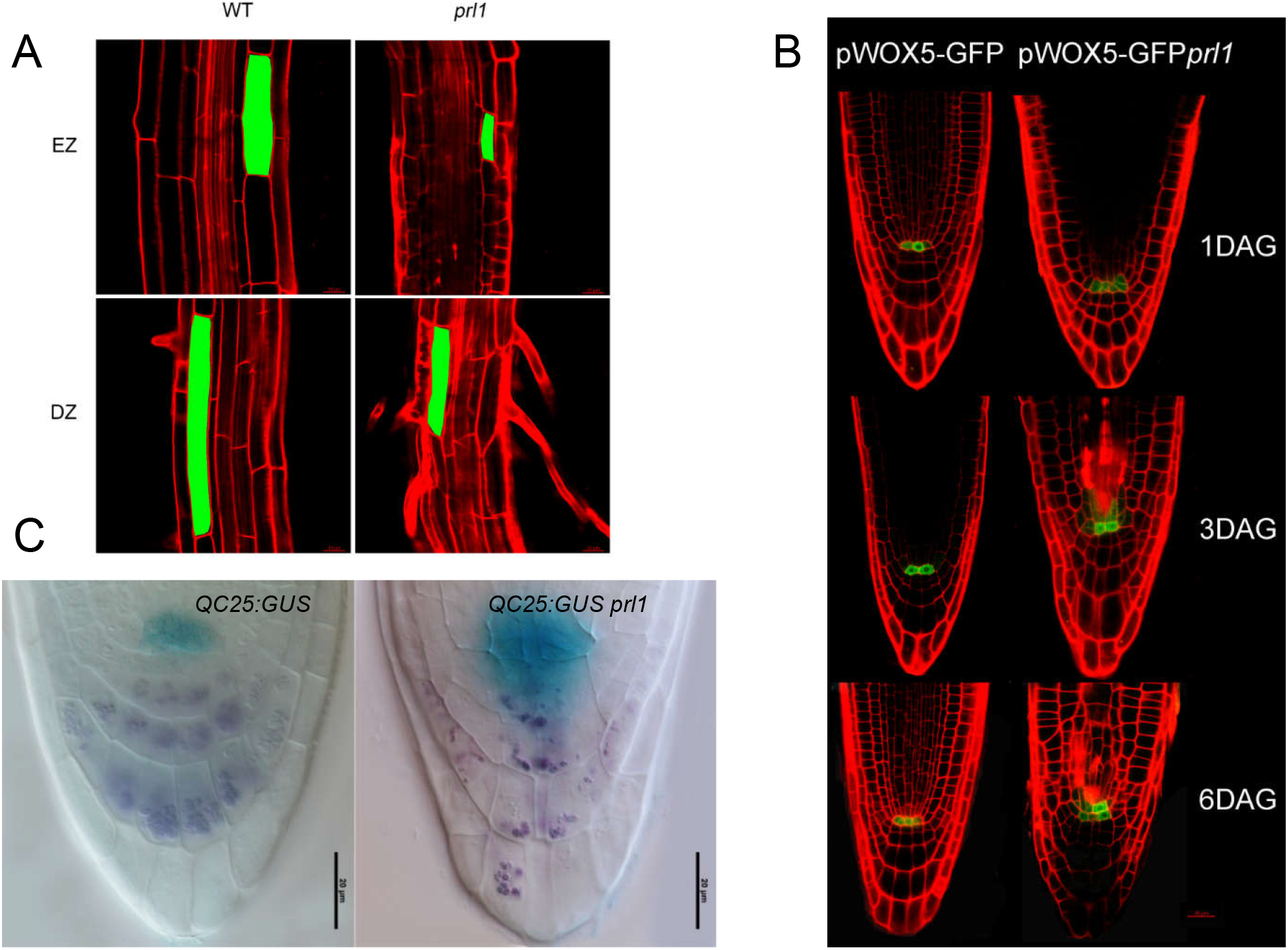
Characters of root tip cells. (A) Cell lengths in the elongation zones and differential zones of WT and the *prl1*. Green color marks a whole cell. Scale bar = 20μm. (B) Time-span pWOX5:GFP distribution and stele cell death in root tips of WT and *prl1*.(C) Distribution of QC25:GUS in root of 6-DAG WT and *prl1*.Bar=20μm.

**Figure S2.**
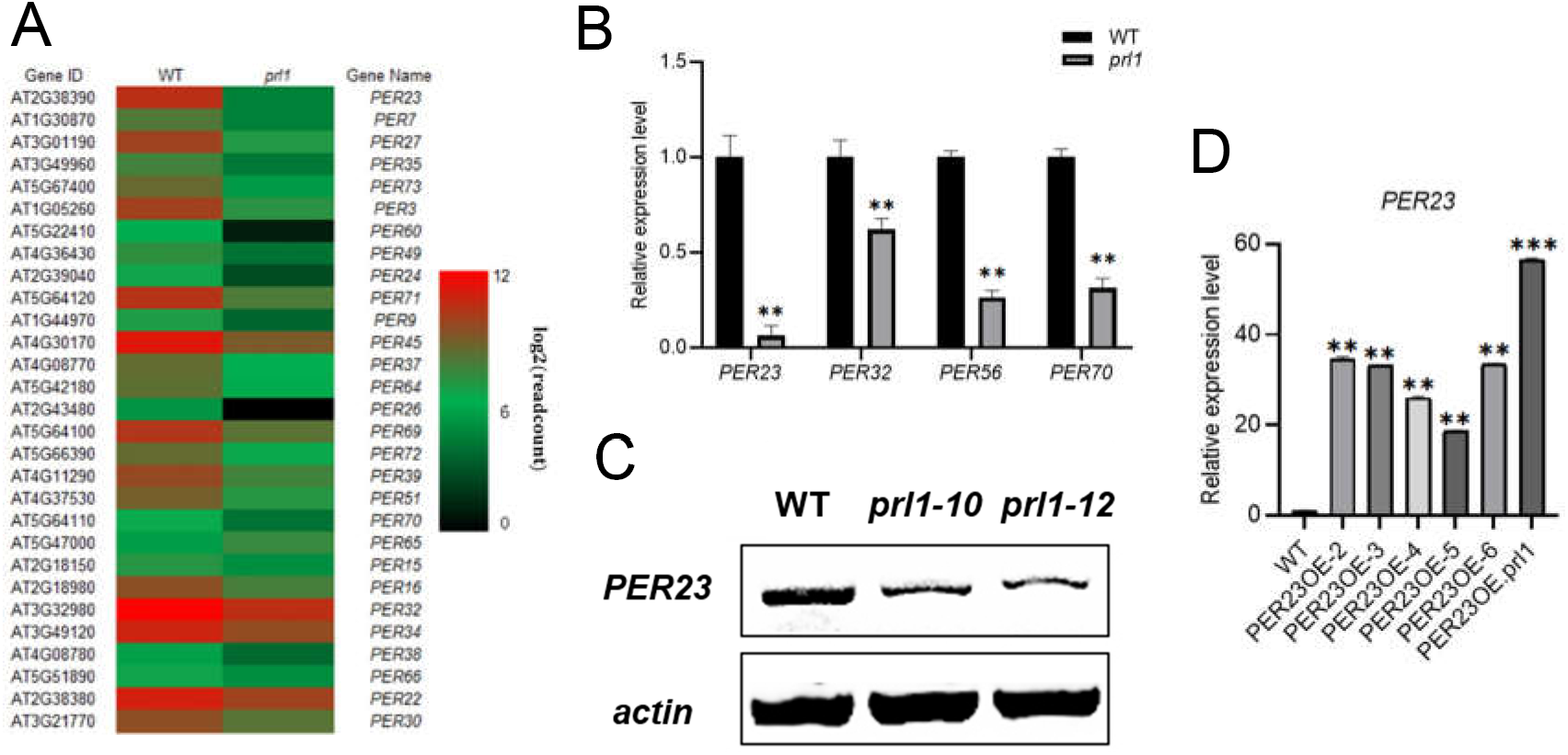
ROS imbalance is undetermined the necessary cue for shortening root of *prl1*. (A) Down expression of peroxidase-encoding genes (*PERs*) in the *prl1*. (B) Relative expression levels of *PER23*, *PER32*, *PER56* and *PER70* in WT and *prl1*. In WT, the expression levels were normalized into “1”. (C) Reverse transcription PCR indicates down-regulation of *PER23* in WT and two allelic lines of *prl1*. (D) Relative expression levels of PER23 in in WT, *PER23* OE lines and *PER23 prl1*. In WT, the expression level was normalized into “1”.

**Figure S3.**
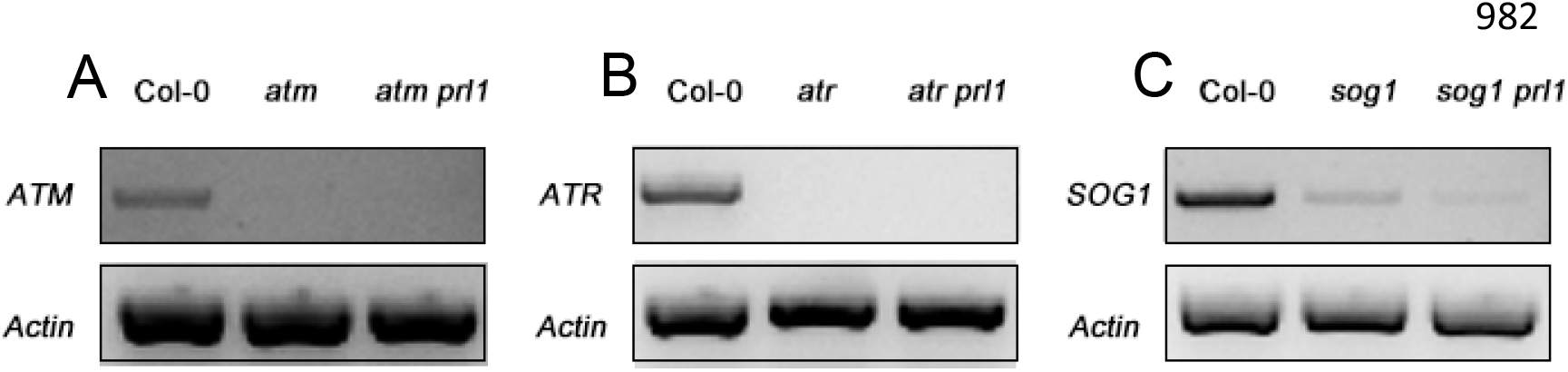
The expression levels of *ATM*, *ATR* and *SOG1* in different genetic backgrounds.

**Figure S4.**
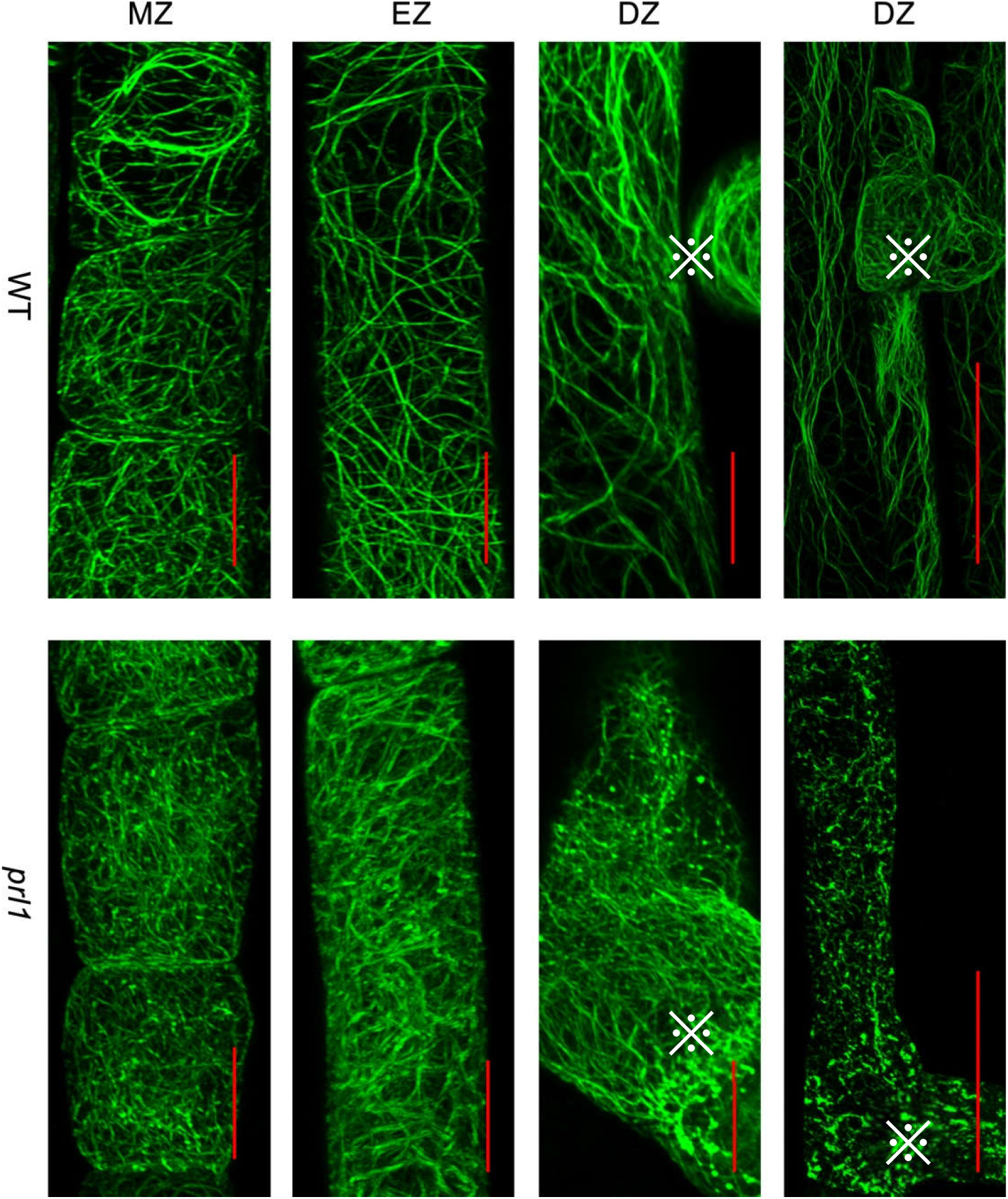
Comparison of cortical actin structures in typical counterpart zones of WT and *prl1*.In three zones of WT, cells succeed in forming filamentous structure; whereas in prl1, actin is disassembled. “※” marks sites of root hairs. Scale bar = 10μm.

**Figure S5.**
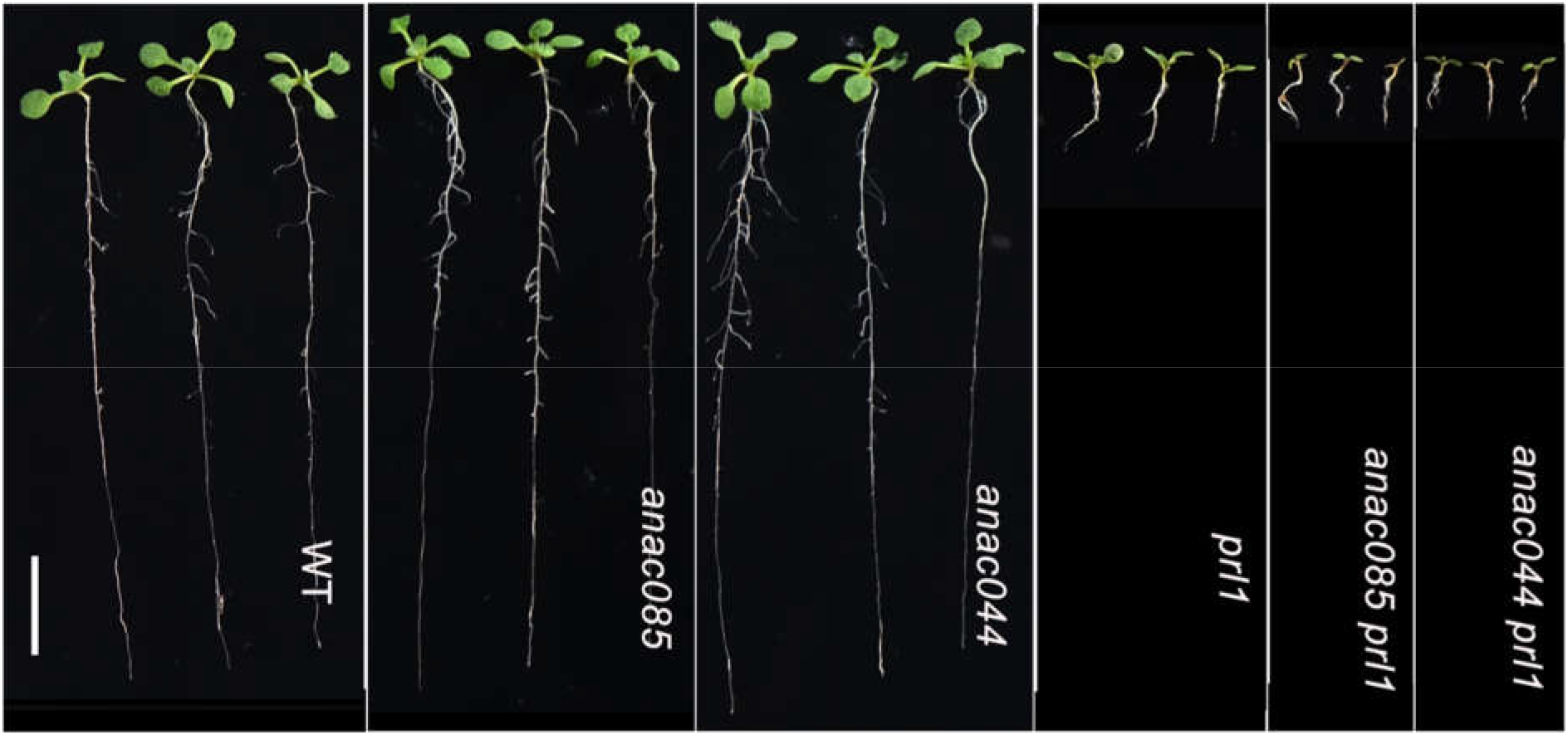
11-DAG seedlings of WT, *anac085*, *anac044*, *prl1*, *anac085 prl1* and *anac044 prl1*.scale bar = 1cm.

**Figure S6.**
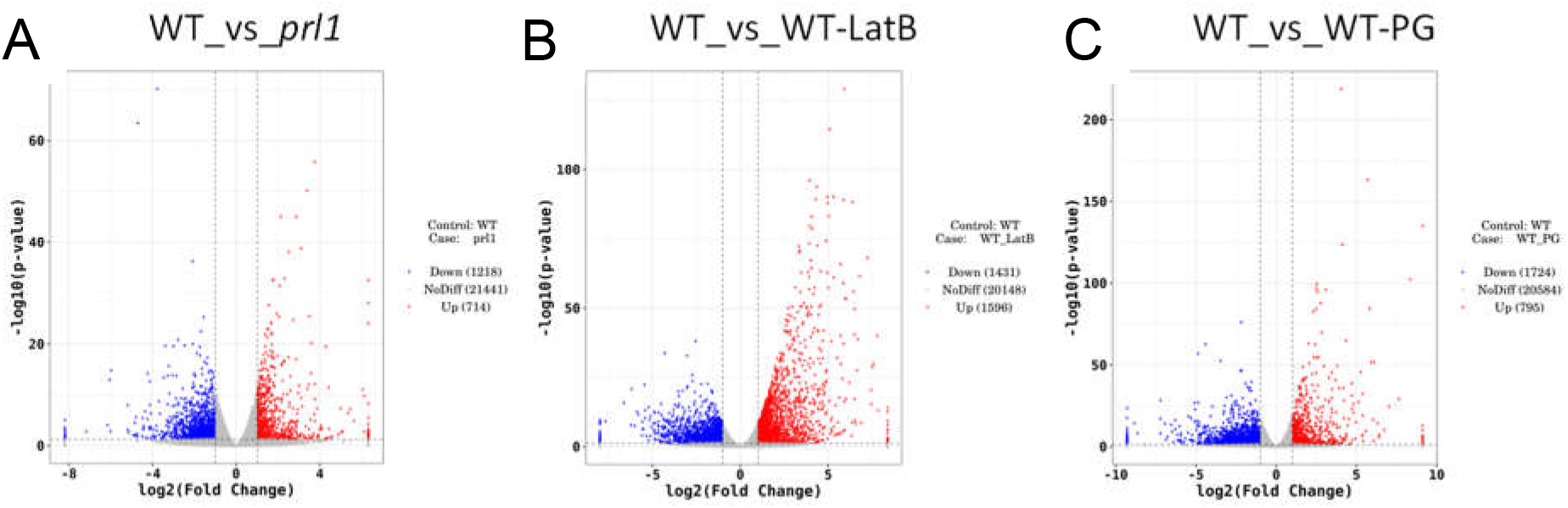
(A) Volcano plot showing fold change levels of the differentially expressed genes (DEGs) *prl1* compared with WT. Significant thresholds for fold change with |log2FoldChange|≥1 and padj≤0.05 are shown in the plot as gray dashed lines. (B) Volcano plot showing fold change levels the differentially expressed genes (DEGs)in LatB-treated WT compared with WT. Significant thresholds for fold change with |log2FoldChange|≥1 and padj≤0.05 are shown in the plot as gray dashed lines. (C) Volcano plot showing fold change levels of the differentially expressed genes (DEGs) in the PG-treated WT compared with WT. Significant thresholds for fold change with |log2FoldChange|≥1 and padj≤0.05 are shown in the plot as gray dashed lines.

### Supplementary tables

**Table S1.**
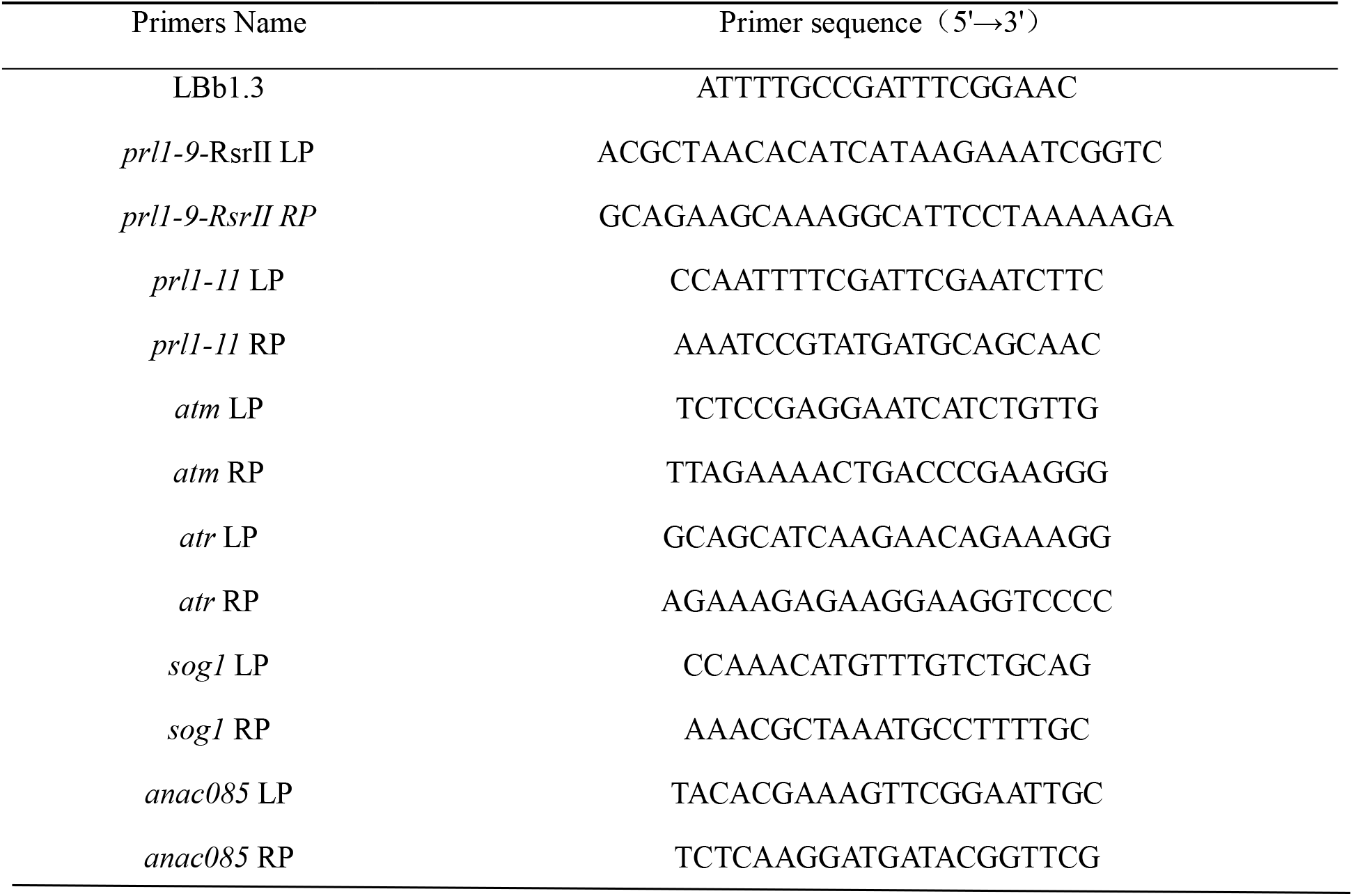
primers used for genotype identification.

**Table S2.**
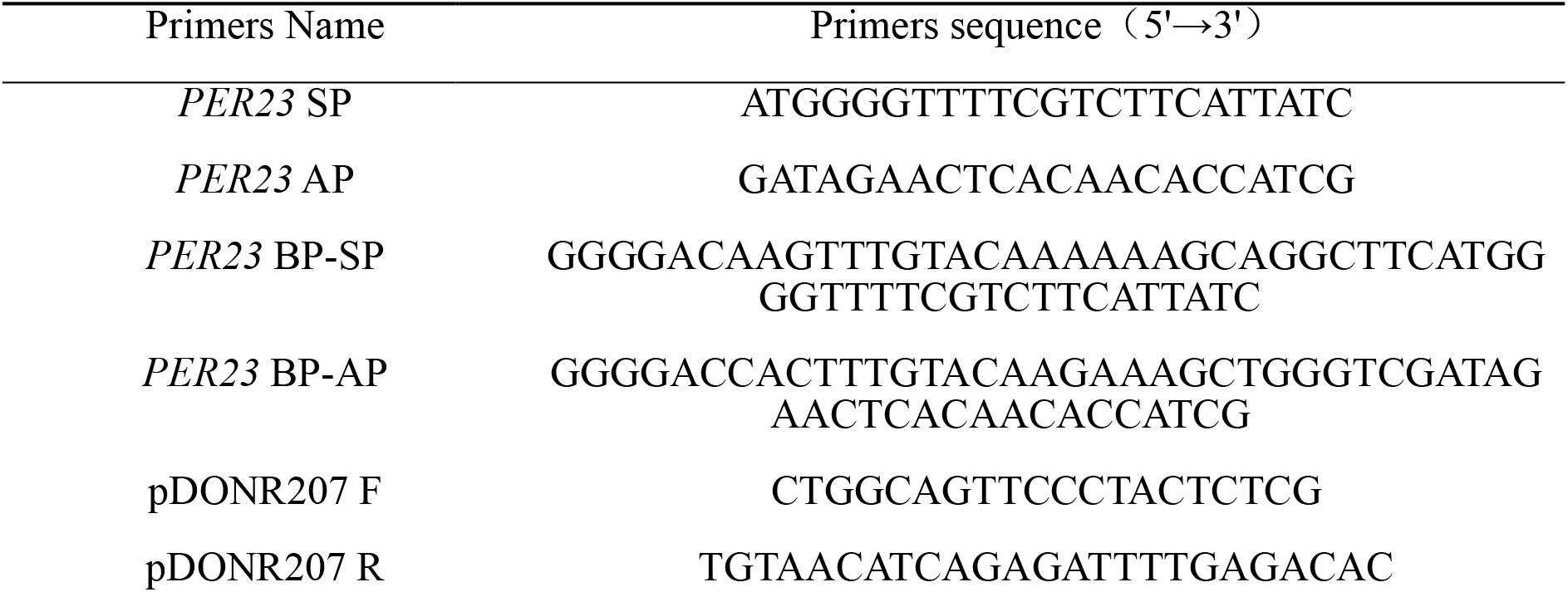
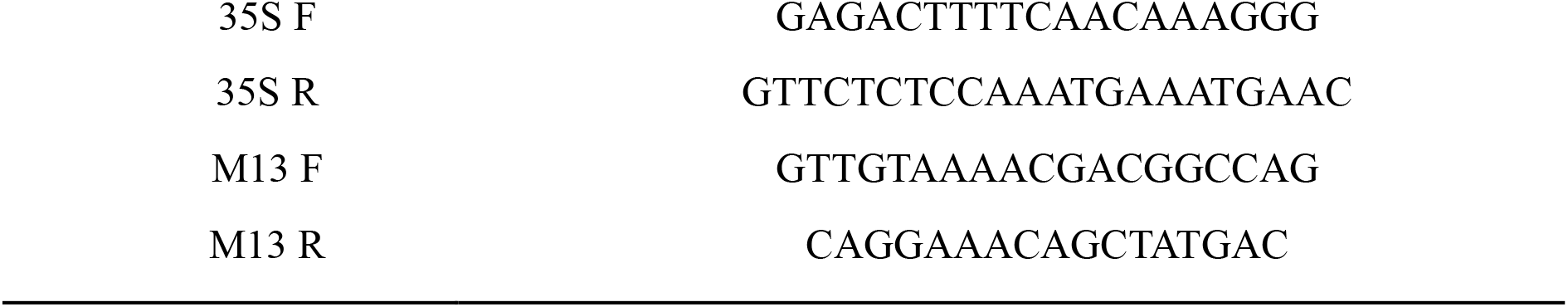
primers used for the construction of *PER23* overexpressed lines.

**Table S3.**
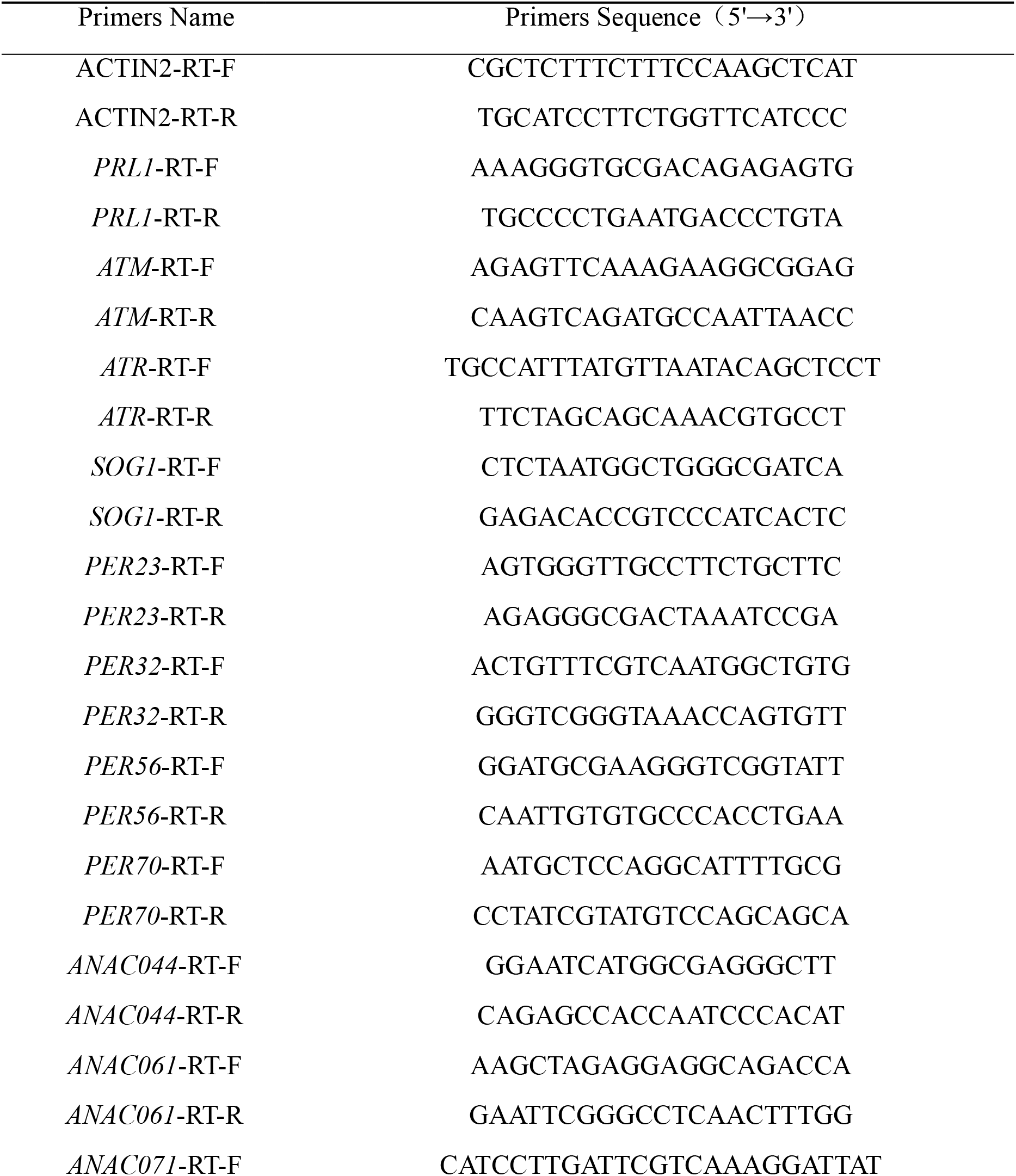
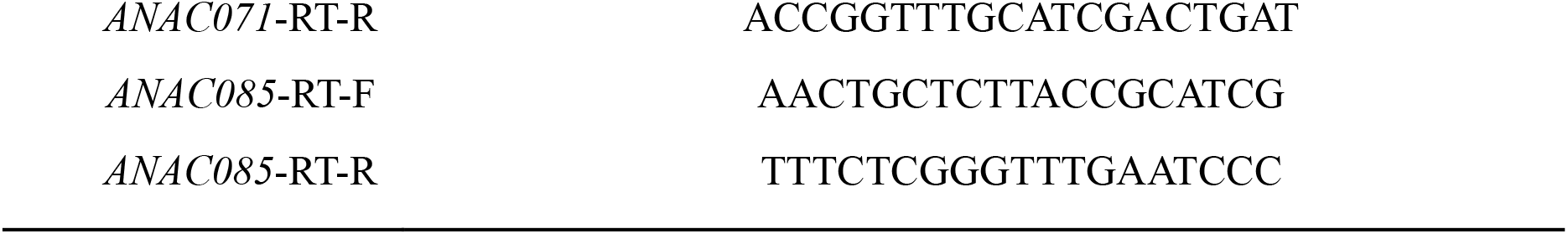
primers used for real-time fluorescence quantitative PCR.

### Dataset S1-S4

